# Familial ALS/FTD-associated RNA-Binding deficient TDP-43 mutants cause neuronal and synaptic transcript dysregulation *in vitro*

**DOI:** 10.1101/2025.03.26.645507

**Authors:** Molly Magarotto, Richard T Gawne, Gabriele Vilkaite, Marcello Beltrami, Andrew S Mason, Han-Jou Chen

## Abstract

TDP-43 is an RNA-binding protein constituting the pathological inclusions observed in ∼95% of ALS and ∼50% of FTD patients. In ALS and FTD, TDP-43 mislocalises to the cytoplasm and forms insoluble, hyperphosphorylated and ubiquitinated aggregates that enhance cytotoxicity and contribute to neurodegeneration. Despite its primary role as an RNA/DNA-binding protein, how RNA-binding deficiencies contribute to disease onset and progression are little understood. Among many identified familial mutations in TDP-43 causing ALS/FTD, only two mutations cause an RNA-binding deficiency, K181E and K263E. In this study, we used CRISPR/Cas9 to knock-in the two disease-linked RNA-binding deficient mutations in SH-SY5Y cells, generating both homozygous and heterozygous versions of the mutant TDP-43 to investigate TDP-43-mediated neuronal disruption.

Significant changes were identified in the transcriptomic profiles of these cells, in particular, between K181E homozygous and heterozygous cells, with the most affected genes involved in neuronal differentiation and synaptic pathways. This result was validated in cell studies where the neuronal differentiation efficiency and neurite morphology were compromised in TDP-43 cells compared to unmodified control. Interestingly, divergent neuronal regulation was observed in K181E-TDP-43 homozygous and heterozygous cells, suggesting a more complex signalling network associated with TDP-43 genotypes and expression level which warrants further study. Overall, our data using cell models expressing the ALS/FTD disease-causing RNA-binding deficient TDP-43 mutations at endogenous levels show a robust impact on transcriptomic profiles at the whole gene and transcript isoform level that compromise neuronal differentiation and processing, providing further insights on TDP-43-mediated neurodegeneration.

## Introduction

TDP-43 is a prominent protein critically involved in neurodegenerative conditions including Amyotrophic Lateral Sclerosis (ALS), Frontotemporal Dementia (FTD) and Alzheimer’s Disorder (AD). It constitutes the pathological inclusions, known as TDP-43 proteinopathy, observed in ∼95% of ALS, ∼50% of FTD (1), and in severe forms of AD (2,3). As a DNA/RNA-binding protein, TDP- 43 is a key regulator of RNA metabolism and processes such as transcription and splicing, and the maintenance and transport of mRNAs (4,5). In disease, TDP-43 mislocalises to the cytoplasm and forms insoluble, hyperphosphorylated and ubiquitinated aggregates (1,6) contributing to enhanced cytotoxicity that is directly linked to neurodegeneration and disease development (1,7,8). In addition to the cytotoxicity directly caused by the aggregates, the sequestering of TDP-43 in aggregates also reduces the levels of functional soluble TDP-43 in the cell, impacting the expression of target genes causing a cascade effect that eventually compromises neuronal health (9,10).

Despite TDP-43’s role as an RNA-binding protein, RNA-binding deficits and RNA dysregulationin disease has been largely overlooked due to hot spots of disease-linked TDP-43 mutations in the aggregation-prone C-terminal of the protein, away from the RNA recognition motifs (RRMs) (11). Currently, TDP-43 is known to directly bind to around 6000 pre-mRNAs in the nervous system, regulating their expression and splicing (4,12). Some of the known targets of TDP-43 such as *STMN2* and *UNC13A* are known to contribute to ALS pathology when misregulated (13–16). The reduction of the full-length *STMN2* and induction of truncated *STMN2* caused by TDP-43 depletion leads to reduced neurite complexity and motor neuron health (15,17). Reduction of *UNC13A* expression causes disruptions in synapse function, contributing to ALS/FTD pathology (13,16). These studies demonstrate that TDP-43 plays a pivotal role in maintaining neuronal health via orchestration of a large network of downstream targets. Many studies have so far employed TDP-43 knockdown as a model. While knockdowns can mimic TDP-43 loss of function, global protein loss does not fully recapitulate the complexity of the disease genetics and pathology.

To further investigate the impact of TDP-43 RNA dysregulation, specifically RNA-binding by TDP- 43 in a disease-relevant model, we focus on two disease-associated TDP-43 mutations, K181E and K263E. Both mutations are identified in ALS and FTD-affected families, linked to the disease phenotypes in an autosomal dominant inheritance pattern (10,18). These mutations are located adjacent to the RNA recognition motifs (RRMs) of TDP-43 and disrupt the binding of TDP-43 to target RNA (10), causing TDP-43 protein to form highly insoluble and phosphorylated nuclear aggregates (10,19). RNA-binding deficient TDP-43-induced nuclear aggregates can sequester endogenous wild-type TDP-43 (10) and potentially other proteins and RNAs, driving a decline in cellular health.

In this study, we used CRISPR/Cas9 to introduce point mutations K181E and K263E into the *TARDBP* gene of human neuroblastoma cell line, SH-SY5Y. Clones of cells carrying either heterozygous and homozygous mutations were generated, which showed the expected change in the splicing pattern of TDP-43 target RNA *PolDip3* as seen in previous studies (10,20). Transcriptomic comparison revealed significant gene expression differences in neuronal and synaptic genes between K181E-TDP-43 homozygous and heterozygous cells compared to wild-type control. Interestingly, many of these genes are up-regulated in K181E-TDP heterozygous cells but down-regulated in K181E-TDP homozygous cells. This finding is validated functionally in the observation of differing degrees of efficiency in neuronal differentiation and neurite growth in the cells carrying the disease-linked RNA-binding deficient mutant TDP-43 with heterozygotes pre-disposed to differentiate and homozygotes unable to differentiate.

## Results

### Establishment and validation of the CRISPR/Cas9-K181E and K263E knock-in SH-SY5Y cell lines

To investigate disease-causing RNA-binding deficient TDP-43-mediated pathogenesis mechanisms, K181E and K263E mutations were introduced to the *TARDBP* gene in human neuroblastoma cell line SH-SY5Y via CRISPR/Cas9 editing (Figure 1A). Potential off-target effects were minimised by using guideRNAs (Figure 1A) with low off-target scores according to ThermoFisher, IDT and Wellcome Sanger Insitute Genome Editing (WGE). From two independent transfections and screening of 110- 130 clones per genotype, multiple clones of either homozygous or heterozygous K181E- and K263E- TDP-43 cells were isolated and validated. The nucleotide substitutions were validated by sequencing the genomic DNA of the targeted area and *TARDBP* cDNA (Supplementary Figure 1A). The integrity of *TARDBP* cDNA was also checked, and no chromosome rearrangement or disruption of *TARDBP* gene was observed (Supplementary Figure 1B).

**Figure 1:**
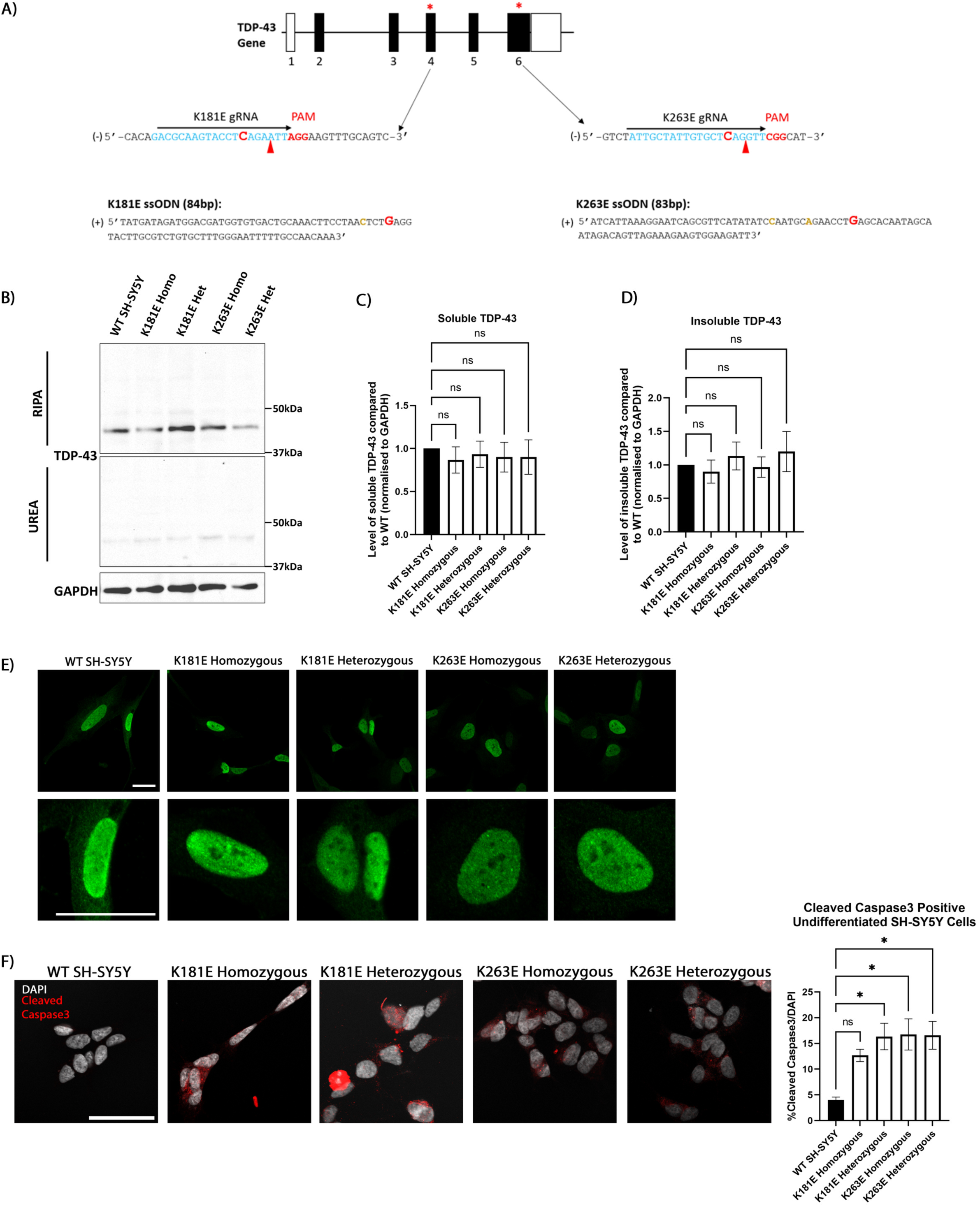
Generating CRISPR-edited K181E or K263E-TDP-43 cells. **A.** The design to introduce K181E and K263E substitution via CRIPSR-editing. Nucleotides encoding K181 and K263 are localised in exon 4 and 6 respectively on TARDBP gene. The position of gRNA, PAM sequence and expected Cas9 cut sites are shown. The gRNAs are highlighted and shown in blue, within which the nucleotide changes for K181E and K263E are highlighted in red. The PAM sequence (AGG or CGG) is immediately downstream of the gRNA alignment site, the expected cut site is indicated with the red arrow head. The sequence used of ssODN is also displayed, with the nucleotide changes for K181E and K263E highlighted in red, and silence mutations in gold. **B.** Immunoblot of K181E and K263E cells showing soluble RIPA and insoluble UREA fractions. TDP-43 antibody used to stain endogenous levels of TDP-43, and GAPDH used as a loading control. **C.** Quantification of soluble TDP-43 levels, normalised to GAPDH. The data were analysed by One-way ANOVA followed by Dunnet’s post-test. **D.** Quantification of insoluble TDP-43 levels, normalised to GAPDH. The data were analysed by One-way ANOVA followed by Dunnet’s post-test. **E.** Staining of TDP-43 of unmodified WT or CRISPR/Cas 9- edited mutant SH-SY5Y cells. Images taken on Zeiss LSM 880 confocal using 63X oil lens. Scale bar = 25µm. **F.** Staining of cleaved caspase-3 of unmodified WT or CRISPR/Cas9-edited mutant SH-SY5Y cells. Scale bar = 50µm.Quantification of cells with cleaved caspase-3 staining (%). 100 cells were counted per condition per experiment, N=3-5. Means and SDs are shown. The data was analysed by One-way ANOVA followed by Dunnet’s post-test. ns = not significant, * p<0.05.

The knock-ins were then functionally validated via examination of the splicing pattern for the most well-known target of TDP-43, *PolDip3* (21), commonly used as a reporter for TDP-43-mediated RNA regulation. Indeed, splicing analysis showed significant reduction of the shorter splicing form of *PolDip3* compared to WT SH-SY5Y cells (Supplementary Figure 1C), consistent with previous reporting that both K181E- and K263E-TDP-43 fail to bind and mediate the exclusion of the cryptic exon 3 of *PolDip3* mRNA (10). Introduction of these disease-linked RNA-binding mutations do not cause the formation of insoluble aggregation of TDP-43 protein or alter its levels (Figure 1B-E) as seen in the case of overexpression (10), despite some nuclear puncta of TDP-43 protein distribution can be observed occasionally (Figure 1E). However, introduction of either K181E or K263E mutations do increase cell death as shown through cleaved caspase-3 staining compared to unmodified cells (Figure 1F).

Overall, we established and validated cell lines using CRISPR/Cas9 to mediate the nucleotide substitution, introducing the disease-causing K181E- or K263E-TDP-43 mutation in either homozygous or heterozygous form where the mutant and wild-type protein is expressed at endogenous level. We validated the cells as disease cell models, observing disrupted splicing patterns of known TDP-43 target mRNA, altered TDP-43 protein distribution and reduced viability.

### Differences in K181E-TDP-43 homozygous and heterozygous transcriptomic profiles in comparison to wild-type cells observed by RNA-seq analysis

To investigate potential cellular dysfunction caused by both K181E and K263E mutations, the transcriptomic profiles of K181E- and K263E-TDP-43 cell lines were established. Bulk RNA-seq was conducted on individual K181E- and K263E-TDP-43 homozygous and heterozygous lines, compared with un-edited SH-SY5Y cells. When comparing K181E-TDP-43 cells with control, 233 and 167 differentially expressed genes (DEGs) were identified in K181E-TDP-43 homozygous and heterozygous cells respectively (q<0.1; Figure 2A-B, Supplementary Data 1). Interestingly, the gene expression changes in K181E-TDP-43 heterozygous versus homozygous cells do not follow an incremental pattern of increased or decreased abundance, nor do these changes occur in the expected direction consistent with mutation dosage. As shown in Figure 2C, the DEGs that are up-regulated in K181E-TDP-43 homozygous cells are often seen down-regulated in K181E-TDP-43 heterozygous, and the genes down-regulated in homozygous are seen up-regulated in the heterozygous cells. Further comparison between K181E-TDP-43 homozygous and heterozygous cells identified 997 genes that express at significantly different levels between the two and at a more stringent significance cut-off (q<0.05; Figure 2D, Supplementary Data 1), highlighting that homozygous and heterozygous expression of these mutant-TDP-43 have divergent impacts on cells. For K263E-TDP-43 cells, no DEGs were identified when comparing K263E-TDP-43 homozygous to control cells, and only 43 DEGs for K263E-TDP-43 heterozygous (q<0.1; Supplementary Figure 2A-B). Interestingly, there was again polarised expression between homozygous and heterozygous observed at a more stringent significance cut-off (q<0.05; Supplementary Figure 2C-D). However, this occurred at a much subtler level overall than K181E. RNA-seq data also showed no evidence of WGE putative off-target exonic editing (Supplementary Figure 3A-C), and no differential expression of genes with potential intronic or downstream of intergenic off-targets (Supplementary Data 2). We also sequenced the genomic area of the top intronic/intergenic off-targets according to WGE, and found no sequence alteration or in/dels (Supplementary Figure 3D-G), suggesting the impact is unlikely to be caused by a CRISPR off-target effect.

**Figure 2:**
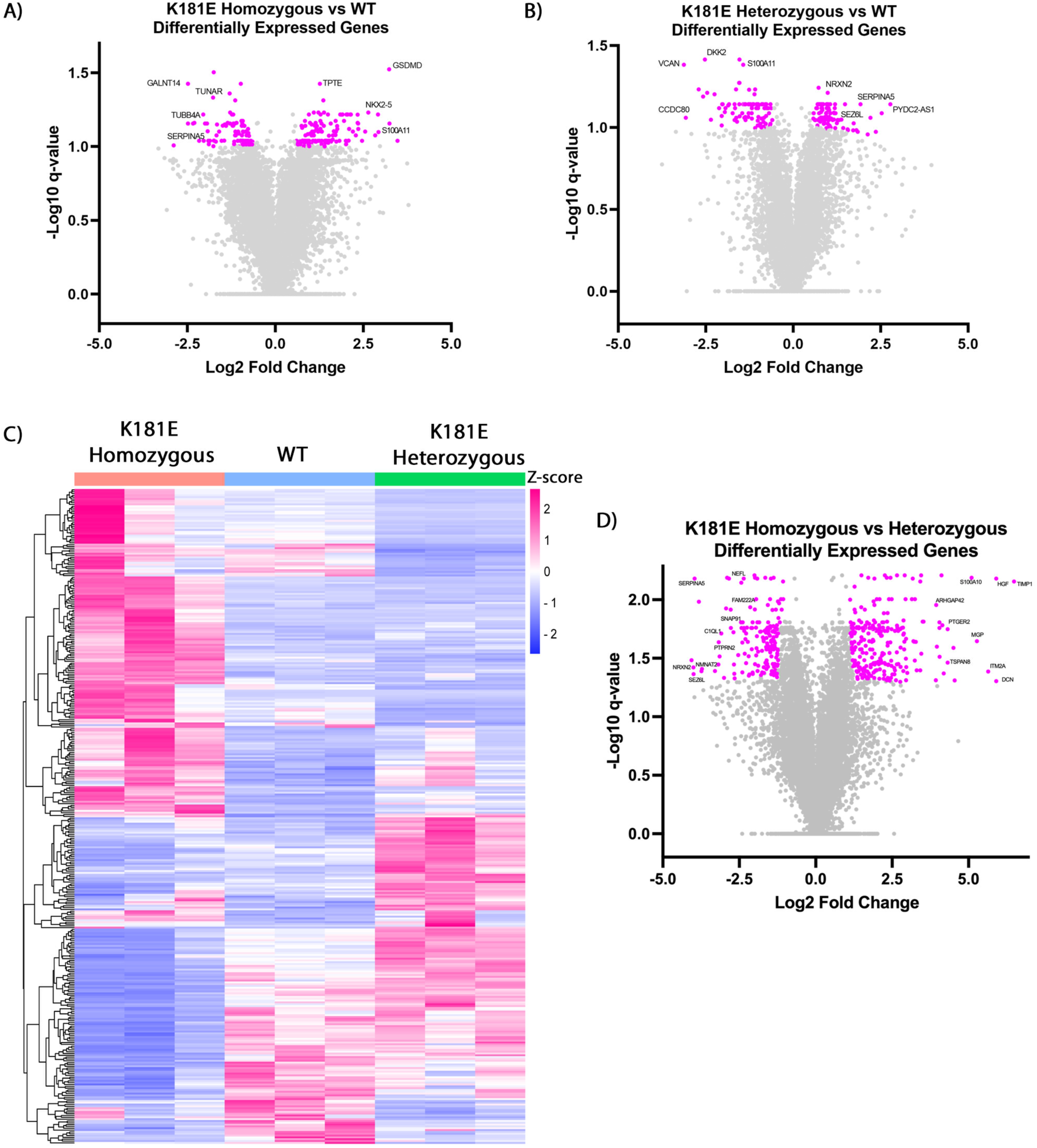
Polarised DEGs in K181E-TDP-43 homozygous and heterozygous cells. **A-B.** Volcano plots showing genes with significantly altered expression in K181E-TDP-43 homozygous cells (A) or heterozygous cells (B) relative to un-edited control SH-SY5Y cells. DEGs highlighted in pink were identified with a cut-off of 0.1 for adjusted *p* values (- Log10(q<0.1)) and a total cut-off of 1 for the log2 fold change ratio. **C.** Heatmap comparing the DEGs level across conditions. All DEGs identified in K181E-TDP-43 homozygous vs WT(B) and K181E-TDP-43 heterozygous vs WT (C) are plotted in the heatmap. The colour scale shows transcripts that are upregulated (pink) or downregulated (blue) relative to mean expression of all samples (Z-score). **D.** Volcano plot showing genes with significantly altered expression in K181E homozygous compared to heterozygous cells. DEGs highlighted in pink were identified with a cut-off of 0.05 for adjusted *p* values (-Log10(q<0.05)) and a total cut-off of 1 for the log2 fold change ratio.

As both mutations disrupt a critical, identified function of TDP-43 protein in interrupting its binding to RNA, it has been suggested that they are loss-of-function mutations for TDP-43 (10,18,20). To examine the hypothesis, we compared the transcriptomic data of our CRISPR/Cas9 knock-in cells with a TDP-43 knockdown dataset (14). No obvious patterns of similarities or differences were observed in Log2 Fold Change (Supplementary Figure 4A) and the datasets had a weak correlation overall (Pearson’s correlation coefficient R = 0.1273; Supplementary Figure 4B). In total, only 12 DEGs were shared in both K181E-TDP-43 homozygous and the TDP-43 knockdown (Supplementary Figure 4C). This observation indicates that while K181E-TDP-43 and K263E-TDP-43 are most noticeable for their disruption in TDP-43 binding to its target RNA, their pathological impact cannot be entirely attributed to the simple loss of function of the TDP-43 protein.

### Neuronal and synaptic pathways are significantly down-regulated in K181E-TDP-43 homozygous cells

To investigate the cellular pathways most affected by K181E mutations, gene set enrichment analysis for gene ontology terms was then carried out to assess DEGs identified in the K181E-TDP-43 homozygous and heterozygous comparison. Due to the low number of DEGs for K263E cells, no significant enrichement for gene ontology terms were identified. Out of the significant pathways highlighted for DEGs down-regulated in K181E-TDP-43 homozygous cells, approximately a third were neuronally related, including those in the top ten most significant (Figure 3A, Supplementary Data 1). Comparing Log2 fold changes to the average transcripts per million (TPMs) for the DEGs in K181E-TDP-43 homozygous showed that the genes most closely associated with the top pathways involving neuronal or synapse growth (Figure 3A) are those showing the greatest negative fold change (Figure 3B). Other significant pathways down-regulated in K181E-TDP-43 homozygous cells included those related to transport, secretion and general biogenesis and signalling (Figure 3C; Supplementary Data 1), again indicating a potential functional compromise in neurite establishment and neuronal communication. In contrast, the gene ontology term analysis of the up-regulated DEGs in K181E-TDP-43 homozygous cells highlighted more generic pathways and functions such as negative regulation of response to stimulus or cell adhesion (Supplementary Figure 5).

**Figure 3:**
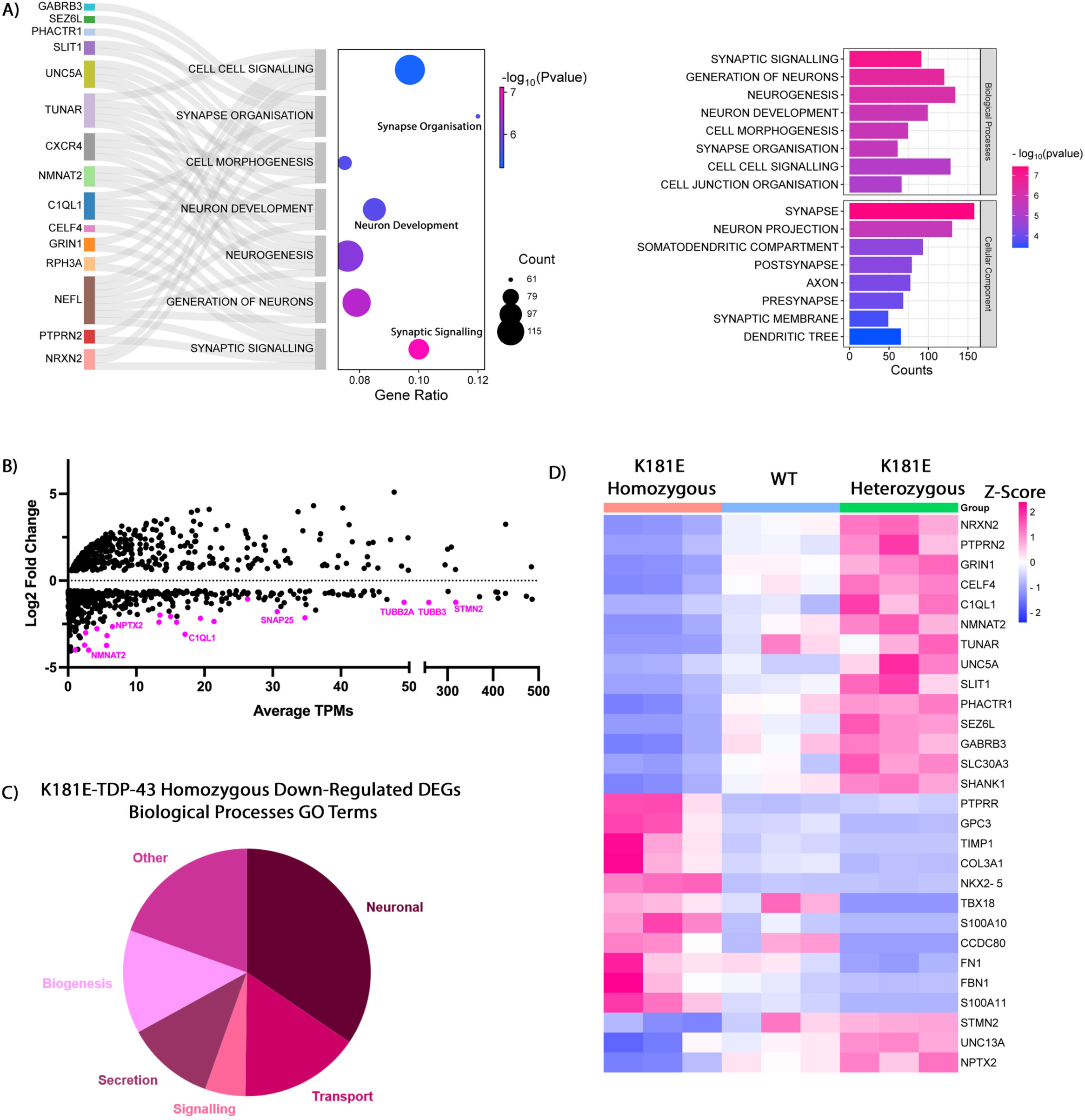
Neuronal and synaptic pathways identified as most significantly down-regulated in K181E-TDP-43 homozygous cells. **A.** Gene set enrichment analysis for DEGs down- regulated in K181E-TDP-43 homozygous cells (compared to K181E-TDP-43 heterozygous), using gene ontology biological processes and cellular components gene sets from MSigDB. πvalues (-log10(p-value) x log2(fold change)) were used as the rank of significance. The most significant gene ontology terms for biological processes and cellular components and associated genes are shown with the associated significance (-log10(Pvalue), shown in colour scale). Gene ratio = (Gene count from RNA-Seq data associated with pathway/ Total gene count within MSigDB pathway dataset). **B.** Log2 Fold Change plotted against Average TPMs for K181E homozygous compared with K181E heterozygous cells. Only DEGs were plotted (q<0.05, total log2 fold change of 1). Genes involved in neuronal pathways identified in gene set enrichment analysis are highlighted in pink. **C.** The frequency of biological processes highlighted in the GO term analysis of genes down-regulated in K181E-TDP-43 homozygous cells. **D.** Heatmap of the most significantly up- and down-regulated genes identified in the K181E-TDP-43 homozygous and heterozygous comparison. The three known TDP-43 targets, STMN2, UNC13A and NPTX2, are also in the comparison.

Notably, the most significant genes linked to neuronal and synaptic functional pathways consistently display an opposing expression pattern in K181E-TDP-43 homozygous compared to heterozygous cells, in that the neuron-related genes are down-regulated in K181E-TDP-43 homozygous cells, but up-regulated in K181E-TDP-43 heterozygous cells compared to unmodified cells, whereas the cellular response genes that are up-regulated in K181E-TDP-43 homozygous cells are reduced in K181E-TDP-43 heterozygous cells (Figure 3D). The known TDP-43 targets implicated in ALS disease pathogenesis, STMN2, UNC13A and NPTX2 (13–16,22), also show this similar opposing expression pattern, being down-regulated in K181E-TDP-43 homozygous cells but up-regulated in K181E-TDP-43 heterozygous cells (Figure 3D).

Overall, our data suggest that K181E-TDP-43 expression significantly compromises the transcriptomic profile of the cells, most significantly the genes involved in neuronal and synaptic functions. The direction of the impact appears to depend on whether the mutant TDP-43 is expressed in the homozygous or heterozygous manner.

### Transcript isoform switching identified between K181E homozygous and heterozygous cells

Following the investigation of changes to gene levels, we next examined whether the expression of K181E-TDP-43 alters transcript isoform shifts which can occur without an overall gene change (23–25). The isoform switching analysis identified approximately 7000 significant isoform changes for K181E-TDP-43 homozygous cells and only around 100 for K181E-TDP-43 heterozygous cells compared to WT cells (q<0.05; Figure 4A-B, Supplementary Data 1), indicating that the homozygous expression of K181E-TDP-43 has a stronger influence on alternative splicing compared to the heterozygous expression, despite a similar number of overall differentially expressed genes. Further analysis found that the types of alternative splicing events were induced to similar levels in both homozygous and heterozygous K181E-TDP-43 (Figure 4C).

**Figure 4:**
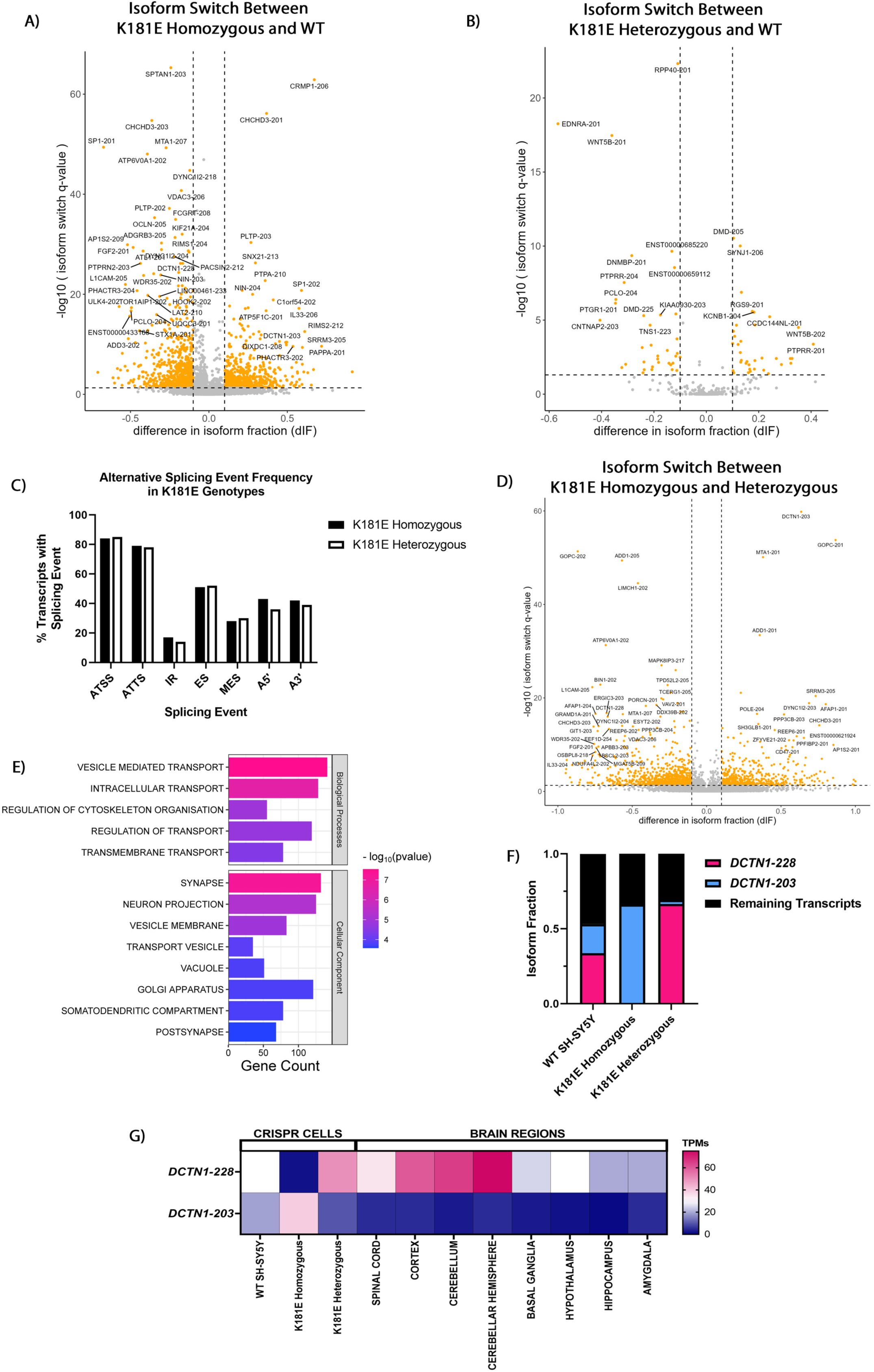
Isoform changes in neuronal genes as a result of K181E-TDP-43 mutations. **A-B.** Volcano plots generated from IsoformSwitchAnalyzeR showing significant isoform switches in K181E-TDP-43 homozygous cells (A) or heterozygous cells (B) relative to un-edited control SH-SY5Y cells. -Log10(q<0.05) plotted against difference in isoform fraction. Significant changes are recognised by an absolute difference in isoform fraction of 0.1, highlighted in yellow. **C.** Alternative splicing events caused by K181E-TDP-43 expression. The alternative splicing events are identified by IsoformSwitchAnalyzeR, analysing all transcript isoforms identified as significantly changed in (A) and (B). ATSS, Alternative Transcript Start Site; ATTS, Alternative Transcript Termination Site; IR, Intron Retention; ES, Exon Skipping; MES, Mutually Exclusive Exons; A5’, Alternative 5’ Splicing; A3’, Alternative 3’ Splicing. **D.** Volcano plot generated from IsoformSwitchAnalyzeR showing significant isoform switches between K181E-TDP-43 homozygous cells and heterozygous cells. Significant changes are recognised by an absolute difference in isoform fraction of 0.1, highlighted in yellow. **E.** Gene set enrichment analysis for significantly changed genes identified in (D), using gene ontology biological processes and cellular components gene sets from MSigDB. Gene set enrichment analysis used πvalues (-log10(p-value) x log2(fold change)) as the rank of significance. The most significant gene ontology terms for biological processes and cellular components and associated gene are shown with associated significance (shown in colour scale). Gene ratio= (Gene count from RNA-Seq dataset associated with pathway/ Total gene count within MSigDB pathway dataset). **F.** Shift of DCTN1 isoforms present in WT, K181E-TDP-43 homozygous and heterozygous cells. The fraction of *DCTN1-228, DCTN1-203* and other DCTN1 isoforms found in our RNA-seq dataset are plotted. Combined fractions of the remaining 26 *DCTN1* isoforms are illustrated in black. **G.** Heat map of gene expression reads (TPMs) for *DCTN1-228* and *DCTN1-203* from various brain regions obtained from GTex (https://gtexportal.org/home/), and our WT and K181E-TDP-43 datasets. Increased abundance indicated in pink, decreased abundance indicated in blue.

Comparing K181E-TDP-43 homozygous to K181E-TDP-43 heterozygous led to greater abundance of genes with isoform switches, totalling around 10,000 significant transcript changes (q<0.05; Figure 4D, Supplementary Data 1), again highlighting the greatest disparity occurring between the two genotypes. This analysis pulled out a novel set of genes compared to the DEGs identified through whole-gene level analysis (Figure 2). However, when applied to the same gene ontology analysis, these genes are again related to neuronal and synaptic pathways, with an enrichment around cytoskeleton and transport (Figure 4E). One of the most significantly affected genes for both K181E- TDP-43 homozygous comparison to WT, and comparison to K181E-TDP-43 heterozygous was *DCTN1*, which encodes dynactin-1, the largest component of the dynactin complex that has critical roles in axonal and cytoplasmic transport, and cytoskeleton assembly (26). We observed a complete isoform switch in K181E-TDP-43 homozygous from the canonical *DCTN1-228* transcript to *DCTN1- 203* (Figure 4F). Using tissue-specific gene expression data acquired from the Genotype-Tissue Expression (GTEx) Portal (https://gtexportal.org/home/), we found that the *DCTN1-228* transcript is highly expressed in several brain regions, while *DCTN1-203* is hardly expressed (Figure 4G). The switch to *DCTN1-203* in the homozygous could contribute functionally to the state of the cells similarly to the overall down-regulation of neuronal genes.

Isoform switching analysis on K263E-TDP-43 homozygous and heterozygous cells in comparison to WT cells only identified around 20 and 100 isoform changes respectively (Supplementary Figure 6A- B; Supplementary Data 1) with similar alternative splicing event frequencies (Supplementary Figure 6C).

In summary, isoform switching analysis highlighted transcript dysregulation caused by changes in splicing events due to K181E-TDP-43 expression. Interestingly, the most prominent changes are again attributed to genes involved in neuronal and synaptic pathways, suggesting the main disruption of K181E-TDP-43 expression would be in neuronal health and functions.

### Neuronal dysregulation caused by K181E-TDP-43

The RNA-seq results suggest K181E-TDP-43 expression most prominently affects the expression and/or transcript isoforms of genes involved in neuronal growth and synapse development (Figure 3-4). To validate this result, we adapted a published protocol to differentiate SH-SY5Y cells (27) into MAP2 and beta III-tubulin positive cells with characteristic neurite projections (Figure 5A). First, the differentiation efficiency was examined using MAP2 as markers of mature neurons. As seen in Figure 5B, reduced MAP2 staining was observed in K181E-TDP-43 homozygous, K263E-TDP-43 homozygous and heterozygous cells across differentiation in comparison to WT cells, but not in K181E-TDP-43 heterozygous cells. In fact, K181E-TDP-43 heterozygous cells had increased MAP2 staining at the undifferentiated stage in comparison to WT. Quantification of MAP2 mRNA (Figure 5C-ii) via RT-PCR was more variable in significance across differentiation than in the staining.

**Figure 5:**
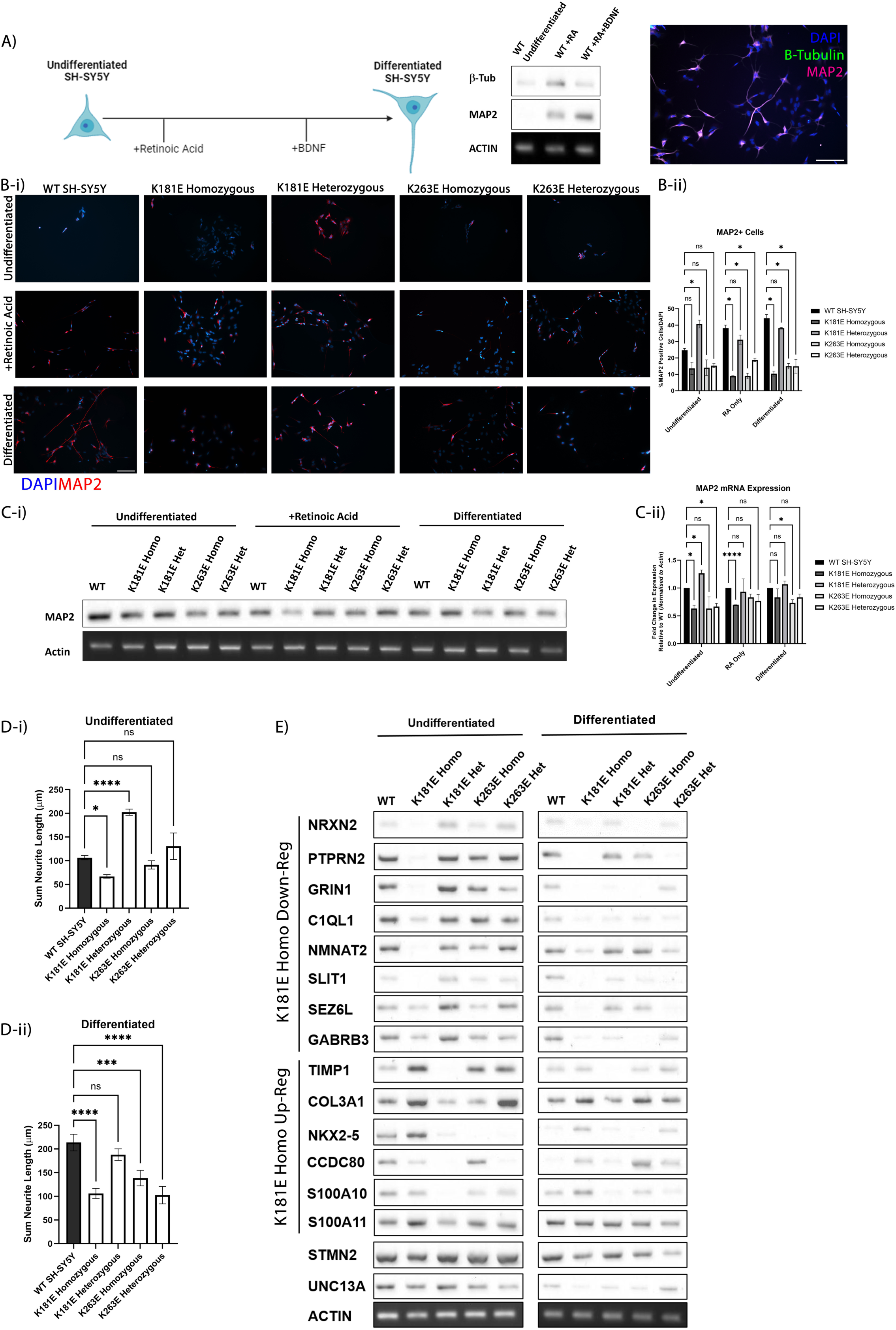
K181E-TDP-43 expression significantly affects neuronal differentiation and processing. **A.** Schematic for SH-SY5Y cell differentiation used in this study (made on BioRender.com). The differentiation is induced via the sequential addition of retinoic acid and BDNF. The RT-PCR and immunofluorescence staining of MAP2 and beta III-tubulin validated the differentiation. **B.** Immunofluorescence (i)showing abundance of MAP2 positive cells in undifferentiated state, mid-differentiation (5 days after retinoic acid treatment) and fully differentiated (5 days after BDNF treatment). The proportion of MAP2 positive cells are quantified from three independent studies (ii). Scale bar = 100µm. Means and SDs are shown. The data were analysed by Two-way ANOVA with Tukey’s post-test. **C.** Expression of MAP2 is visualised and quantified by RT-PCR of endogenous MAP2 mRNA at different differentiation stages (i). RT-PCR of actin is used as loading control. The levels of MAP2 were quantified, normalised to the loading control, and presented in relation to WT (ii). N=3. Means and SDs are shown. The data were analysed by Two-way ANOVA with Tukey’s post-test. **D.** Sum neurite length from MAP2 staining from (B) for undifferentiated (i) and differentiated (ii) conditions were quantified for individual cells across three independent experiments for the individual clones. Means and SDs are shown. The data were analysed by One-way ANOVA followed by Dunnet’s post-test. **E.** Validation of the changes in genes-linked to neuronal development and progression identified in our K181E-TDP-43 RNA seq dataset. RT-PCR of selected genes identified in RNA-seq was carried out using RNA extracted from cells of undifferentiation or fully differentiation state. Known TDP-43 targets, STMN2 and UNC13A are also included in this study. Actin is used as a loading control. A representative RT-PCR result is shown here with the overall quantification and statistically comparison from 3 independent extractions shown in supplementary figure. ns = not significant, * p<0.05, ** p<0.01, *** p<0.001, **** p<0.0001.

However, the trends generally showed that MAP2 mRNA was reduced in mutant cells, except for K181E-TDP-43 heterozygous cells (Figure 5C). This is consistent with the earlier observation in the RNA-seq study that the neuronal-linked genes are down-regulated in K181E-TDP-43 homozygous cells but up-regulated in K181E-TDP-43 heterozygous cells (Figure 3D). From the cells that were positive for MAP2 (Figure 5B-i), the sum neurite lengths of individual cells were subsequently quantified (Figure 5D). As shown in figure 5D, the neurite lengths of the MAP2-positive differentiated cells were reduced in K181E-TDP-43 homozygous cells (Figure 5D-ii). However, K181E-TDP-43 heterozygous cells exhibited longer neurites compared to control in undifferentiated condition (Figure 5D-i) and similar neurite length to control when differentiated (Figure 5D-ii).

Examination of the neuronal-linked genes identified in the RNA-seq confirmed that they were down- regulated in K181E-TDP-43 homozygous cells but up-regulated in K181E-TDP-43 heterozygous cells in the undifferentiated condition (Figure 5E; Supplementary Figure 7A) which could be the reason why K181E-TDP-43 homozygous cells cannot differentiate efficiently and why K181E-TDP-43 heterozygous cells appear to be primed with MAP2 induction and longer neurites even in the undifferentiated condition (Figure 5B-D). The gene expression after differentiation showed a similar pattern, that neuronal-linked genes were still reduced in K181E-TDP-43 homozygous cells (Figure 5E; Supplementary Figure 7B). Their levels in K181E-TDP-43 heterozygous cells were mostly similar to WT, however, some neuronal genes exhibited decreased expression in a similar manner to K181E-TDP-43 homozygous cells, including *PTPRN2*, *GRIN1* and *SLIT1* (Figure 5E; Supplementary Figure 7B). Since K181E-TDP-43 heterozygous differentiated cells displayed similar neurite growth and overall morphology as WT cells once differentiation was initiated, potentially the genes that begin to align with the homozygous state are less vital for these processes to occur.

The examination of selected genes seen up-regulated in K181E-TDP-43 homozygous cells also verified the earlier results from RNA-seq, in that the up-regulation in K181E-TDP-43 homozygous cells and down-regulation in K181E-TDP-43 heterozygous cells was validated in the RT-PCR (Figure 5E; Supplementary Figure 7C). This pattern was largely maintained in differentiated cells except for *S100A11* and *COL3A1* whose expression level seem to even out in all genotypes in differentiated cells (Figure 5E; Supplementary Figure 7D). Finally, the expression levels of the known TDP-43 targets, *STMN2* and *UNC13A*, were also validated in these cells in both undifferentiated and differentiated conditions. A more significant difference was observed between K181E-TDP-43 homozygous and WT cells after differentiation (Figure 5E; Supplementary Figure 7E-F), highlighting them as more mature neuron TDP-43 targets. K181E-TDP-43 heterozygous cells began to see a reduction in *UNC13A* after differentiation, but not in *STMN2* (Figure 5E; Supplementary Figure 7F).

The K263E mutation affects RRM2 which is proposed to play a minor role in binding to TDP-43’s target RNA (5,28) especially when the mutant protein is expressed at endogenous levels. Whereas no prominent transcriptomic profile change was observed using undifferentiated K263E-TDP-43 homozygous and heterozygous cells, they did begin to exhibit some deficits once differentiation begins (Figure 5B-D) with the neuronal-linked genes down-regulated in a similar manner to K181E- TDP-43 homozygous, alongside known targets *STMN2* and *UNC13A* (Figure 5E; Supplementary Figure 7). All functional observations were confirmed in independent lines for K181E-TDP-43 heterozygous and K263E-TDP-43 homozygous and heterozygous obtained in independent CRISPR transfection (Supplementary Figure 8).

## Discussion

TDP-43 pathology is a contributing factor to ∼95% of ALS and ∼50% of FTD (1), and is also strongly associated with severe AD (2,3). Characterisation of the cause-and-effect underlying TDP-43 proteinopathy is therefore a necessary key in unravelling the disease, and to develop targeted therapy. In this study, we developed CRISPR/Cas9 knock-ins of two ALS/FTD-associated RNA-binding deficient TDP-43 mutants, K181E and K263E, into human neuroblastoma SH-SY5Y cells, creating homozygous and heterozygous K181E- and K263E-TDP-43 cell lines that cause splicing changes of known TDP-43 targets such as *Poldip3* (Supplementary Figure 1) (21) and exhibit increased cell death (Figure 1F). Further transcriptomic analysis identified significant changes in K181E-TDP-43 cells, particularly highlighting decreases in expression of genes involved in neuronal and synaptic development, organisation and signalling (Figure 3-4). The functional impact of this transcriptional dysregulation was validated in cell studies where the disruption in neuronal differentiation and neurite growth was observed in all TDP-43 mutant cells except for K181E-TDP-43 heterozygous cells which show hints of being primed for neuronal differentiation even in basal condition (Figure 5).

### TDP-43 RNA-Binding Deficiency and Disease Development

There are well over 1000 RBPs in either the nucleus or the cytoplasm regulating mRNA stability, transport and maturation (29,30). Typically, these proteins contain RNA recognition motifs (RRMs), nuclear import and export signals, and aggregation-prone low complexity domains (LCDs) (31–33). They can form RNA-protein (RNP) granules that can have different roles depending on contributing factors (34–37).RNA regulation and RBPs play pivotal roles in neuronal health and maintenance (38–40), and have been associated with several neurodevelopmental and neurodegenerative disorders (1,3,41–49). One prominent RBP in neurodegenerative diseases is TDP-43, associated with ∼97% of ALS cases, ∼50% FTD cases, and severe cases of Alzheimer’s (1,3,18,50). TDP-43 is ubiquitously expressed, and knock-outs in mice lead to embryonic lethality (51). Overexpression models are also seen to cause cytotoxicity, meaning regulated TDP-43 levels are vital in maintaining cell health (22,52,53).

Despite the close association between TDP-43 proteinopathy and ALS/FTD, the involvement of TDP- 43 RNA dysregulation and binding defects in a disease context havebeen overlooked for a long time due to the hot spots of disease-linked TDP-43 mutations concentrated in the C-terminal aggregation-prone area of the protein, away from the RRMs (11). *In vitro* studies have found that the C-terminal mutations such as Q331K and M337V do not interfere with TDP-43 binding to its target RNA (10), and that the expression of the engineered RNA-binding deficient F147/149L-TDP-43 does not cause any further deficiency in cell function or survival (54,55). The question remains: to what extent is RNA-binding disruption an integrated mechanism of TDP-43 disease-driving?

In 2009 and 2019, the two TDP-43 RNA-binding deficient mutations, K181E and K263E, were identified in independent families of ALS and FTD (10,18). Both mutations cause hyperphosphorylated and insoluble nuclear TDP-43 aggregates in both patient tissue and through overexpression models (10,18), and unlike with the artificial F147/149L-TDP-43, these ALS/FTD-linked mutations lead to cellular disruption and cytotoxicity. As shown in this study, despite no overt nuclear aggregates besides infrequent nuclear puncta observed when the mutant protein was expressed at the endogenous level (Figure 1B-E), due to either the lack of challenge, or the endogenous expression being insufficient to drive aggregation formation, the introduction of these mutations does lead to global transcriptomic changes in gene expression (Figure 2-4) and affect cell survival and function (Figure 1C, 5).

While K181E- and K263E-TDP-43 were observed to have similar affects in the *PolDip3* splicing assay (Supplementary Figure 1C), this was not extended to the RNA-sequencing results themselves. K263E-TDP-43 showed limited differentially expressed genes when compared to unmodified SH- SY5Y cells in their undifferentiated state (Supplementary Figure 2). This is somewhat unsurprising as Imaizumi et al. (20) have previously reported no differences in K263E-TDP-43 homozygous knock-in in undifferentiated iPSCs. Once differentiated, we did observe some significant changes to gene expression through RT-PCR and consequential effects on neuronal health, including a down- regulation of *STMN2* and *UNC13A* (Figure 5; Supplementary Figure 7), consistent with published K263E models and broader knockdown models (13–16,20). On the other hand, K181E-TDP-43 homozygous and heterozygous cells did show more significant changes in transcriptomic profile in comparison to the WT cells (Figure 2). Unlike the K263E adjacent RRM2 which is suggested to play a supporting role in facilitating RRM1-RNA binding (56), K181E is found adjacent to RRM1 which is readily associated to TDP-43’s autoregulation, managing its levels in the nucleus (56,57), and has been associated with TDP-43’s ability to undergo liquid-liquid phase separation during cellular stress (58).

In general, ALS and FTD are age-related diseases where aberrant proteins and harmful substances accumulate over time. Whereas wild-type or LCD mutant TDP-43 can still bind to target RNA, overexpression leads to splicing misregulation, similar to observations in TDP-43 knock-down (10,23,25,59). This is likely due to the changes in the pool of functional TDP-43 via its nuclear clearance and increased aggregation, lending support to the theory that TDP-43 loss-of-function is a driver of pathology. This is further supported by knock-down studies which identified around 100 cryptic exon-containing transcripts which are the targets of nonsense-mediated decay (NMD) and other degradation pathways (60), leading to protein down-regulation. Two prominent ALS-linked targets of TDP-43 are *STMN2* and *UNC13A*. Under TDP-43 depletion, the full-length *STMN2* decreased while the truncated *STMN2* increased through cryptic exon inclusion, leading to deteriorated motor neuron health (14,15,17). *UNC13A* is a critical gene for synapse function, with SNPs near to the cryptic exon of *UNC13A* seen to upregulate cryptic splicing and alter the ability of TDP-43 to bind and repress the cryptic exon inclusion. The resulting transcript is degraded by NMD, decreasing the overall level of the functional protein (13,16). The changes of TDP-43 targets such as *STMN2* and *UNC13A* are also seen in ALS and FTD patient tissues (13–16), and the overexpression of full-length *STMN2* and use of antisense oligonucleotides to rectify *UNC13A* cryptic exon inclusion rescue toxicity in ALS cell models (14,61–63), suggesting that disruption of TDP-43-mediated RNA regulation is indeed a contributing factor for ALS development. Furthermore, high levels of cryptic exon inclusion have also been identified in AD, with *STMN2* and *UNC13A* cryptic exons detected in TDP-43-associated AD (64,65), again, highlighting the role of TDP-43-mediated RNA regulation in neuronal health and the importance in getting a better understanding of its impact. However, it should be noted that the loss-of-function cannot be the sole cause of TDP-43-mediated pathogenesis as no cellular toxicity is seen by the expression of non-disease-linked RNA-binding deficient F147/149L-TDP-43. The dysregulation of RNA-regulation is likely an aggravating factor that jointly with TDP- 43 gain-of-toxicity triggers disease development and progression.

### Dosage-dependent opposing effect of K181E-TDP-43

Most interestingly from the RNA-seq data is that the greatest difference in gene expression occurred between the two K181E-TDP-43 genotypes, where we see the DEG changes in opposite directions in the homozygous and heterozygous K181E-TDP-43 cells, either side of the unmodified wild-type (Figure 2). This polarised gene expression also occurred at a more moderate level in K263E-TDP-43 cells (Supplementary Figure 2). The genes that are down-regulated in the K181E-TDP-43 homozygous and up-regulated in the heterozygous were significantly associated with neuronal and synaptic processes (Figure 3). This feature is observed at both overall gene level (Figure 2) and also on the level of transcript isoform shift (Figure 4).

The down-regulation of these genes or the shift to non-neuronal isoforms in K181E-TDP-43 homozygous cells compromises neuronal differentiation, while opposing expression of these genes in K181E-TDP-43 heterozygous cells appear to prime them for neuronal differentiation in the absence of stimulating factors (Figure 5). This could imply a compensatory mechanism occurring to protect neuronal growth at the detriment to other processes, since these cells exhibit similar cytotoxicity as the other genotypes (Figure 1F). Neuronal overgrowth can occur in response to cellular stress across several models, with this tending to be a temporary compensation of the neurons until the stress is removed (66–70). In disease, overproduction of neurons, dendritic overgrowth and excess synaptic connections have been associated to neurodevelopmental disorders such as autism spectrum disorders (ASD) (71–73), and also with neurodegenerative diseases. A recent study by Capizzi et al. (74) observed that whereas axons from Huntington Disease (HD) mouse model brains were shorter and less able to cross the corpus callosum to the other hemisphere, the axons that were able to cross, had increased dendritic branching, hypothesised by the researchers to be a compensatory action in response to the lower number of surviving neurons. In Parkinson’s Disease (PD), increased neuronal connectivity as a result of PD-related genetic mutations were correlated to severity and aggressiveness of the disease in patients (75). And in ALS, neurite overgrowth and enlarged axonal arbors to compensate the loss of NMJ innervation was observed in mouse models (76,77). This again signifies a balance between pathogenic and compensatory mechanisms in development, with disease-associated factors even at times playing a protective role in early life. The link between neurodevelopmental and neurodegenerative disorders is a tenuous one. In terms of ALS and FTD, several studies have shown their correlation and co-morbidity to neurodevelopmental and psychiatric disorders, often related to psychological and physical hyperactivity (78–80). Cellular stressors have been linked to these neurodevelopmental disorders (81–83), but how this may or may not develop into neurodegenerative diseases is not understood.

Overall, our study demonstrated the impact on transcriptomic profiles by disease-linked TDP-43 RNA-binding deficiency. The most affected pathways are related to neuronal development, synaptic processes and cellular transport, and as such, had functional consequences on the cell’s differentiation and survival. K181E-TDP-43 heterozygous expression appeared to aggravate cellular disturbance in an opposite direction to homozygous K181E-TDP-43 cells. The extent of the disruption correlating to ageing and subsequent challenges still needs to be further investigated, but could potentially shed light on the early-stage pathogenesis for disease.

## Methods and Materials

### CRISPR/Cas9-mediated TDP-43 genome editing via nucleofection

GuideRNA (gRNA) and HDR template (ssODN) (Figure 1A) were designed using the IDT Custom Alt-R^TM^ CRISPR-Cas9 guide RNA and Alt-R HDR Design tools (IDT), where its on-target efficiency and off-target cuts were also estimated. The sequence of crRNAs used in this study are: K181E 5’- GACGCAAGTACCTTAGAATT-3’, K263E: 5’-ATTGCTATTGTGCTTAGGTT-3’. HDR templates used in this study are K181E 5’- TATGATAGATGGACGATGGTGTGACTGCAAACTTCCTAATTCTGAGGTACTTGCGTCTGTGCTTTGGGAATTTTTGCCAACAAA-3’, K263E 5’- ATCATTAAAGGAATCAGCGTTCATATATCCAATGCAGAACCTGAGCACAATAGCAATAGACAGTTAGAAAGAAGTGGAAGATT-3’. 100μM of sequence-specific crRNAs (IDT) were mixed with 100μM of tracrRNA (IDT), heated to 95°C for 5 minutes to form 50μM of gRNA which were subsequently aliquoted and stored at -20°C.

SH-SY5Y cells were cultured using Dulbecco’s modified Eagle medium: Nutrient mixture F12 (DMEM/F12) (Invitrogen) supplemented with 10% foetal bovine serum (Invitrogen), maintained at 37°C, 5% CO^2^ and reached the confluency of 60-80% at the day of transfection. The CRISPR transfection was carried out using nucleofection where 1 million cells resuspended in electroporation buffer (90mM Sodium Phosphate, 5mM KCl, 0.15mM CaCl2, 50mM HEPES) were mixed with the RNP complex (2.5μM gRNA, 4mg/ml polyglycolic acid (PGA) and 1.25μM Alt-R® S.p. HiFi Cas9 Nuclease V3 (IDT)), HDR template (IDT, 2μM) and electroporation enhancer (IDT, 1μM).

Electroporation was carried out using a 2mm cuvette (VWR) and the G-004 programme on the Nucleofector™ II/2b Device (Lonza). Immediately after electroporation, cells were plated in a 6-well plate in culture medium containing 1μM of HDR enhancer (IDT) for 48 hours before clonal selection and genotyping.

### Clonal selection and genotyping

Cells were re-plated in 100 mm dishes at either 1000 or 2500 cells per plate 48 hours after nucleofection and were left to grow into individual colonies which were picked in the next 1-2 weeks. The colonies of cells were then dispersed with trypsin and plated into two identical 96-well plates, one for genotyping and one for further expansion. The cells for genotyping were lysed in lysis buffer (10mM Tris-HCL 7.5pH, 50mM KCl, 1.5mM MgCl2, 0.45% Tween20, 0.5% Triton-X, 20mg/ml Proteinase K). The cell lysates were incubated on the thermocycler for 10 minutes at 56°C to activate the proteinase K followed by 10 minutes at 95°C to stop the enzyme reaction. The crude cell extract was used as template for PCR amplifying fragments containing the CRISPR-edited sites using primers of 5’-GTTAACACAGTATGGATTC-3’ and 5’-TATTCAGCATGCACTAAGG-3’ to generate a 441bp PCR product to genotype K181E site, and 5’-TCGACTGAAATATCACTGC-3’ and 5’- AACCCCACTGTCTACATTCCC-3’ to generate a 691bp PCR product to genotype K263E site. PCR was conducted using the VeriFi™ Polymerase reagents (PCR Biosystems) and ran under recommended thermocycler conditions. PCR products were enzymatically cleaned using ExoSAP- IT™ (ThermoFisher) before submission for Sanger Sequencing (Genewiz, Azenta Life Sciences) using the following sequencing primers: K181E forward 5’-AGCCACTGCATCCAGTTGAAACC- 3’, K181E reverse 5’-TTTCATGAACACACCCTGCCGC-3’, K263E forward 5’- CAGTCTCTTTGTGGAGAGG-3’ and K263E reverse 5’-TCCCTCTGCATGTTGCCTTG-3’.

Primers for potential off-target editing PCR and sequencing are detailed in Supplementary Table 1, in/dels detected using online tool DECODR (84).

### RNA extraction and semi-quantitative RT-PCR

RNA was extracted using the RNeasy® Mini Kit (Qiagen) with DNase I treatment. cDNA was generated using the SuperScript™ III First-Strand Synthesis Kit (Invitrogen) with 50ng/µL random hexamer primer (Thermo Fisher). 50ng of cDNA was used for semi-quantitative RT-PCR. Primers (Table 1) were designed using Ensembl (https://www.ensembl.org/index.html) and ncbi primer blast (https://www.ncbi.nlm.nih.gov/tools/primer-blast/), unless otherwise stated. RT-PCR reactions were conducted using the VeriFi™ Polymerase reagents (PCR biosystems). Tm and the number of cycles used for each gene were detailed in Table 1. All products were run on 2-2.5% agarose gels. The fluorescent tag IRDye 700 was attached to the 5’ end of the forward primer of *Poldip3*, the splicing assay of which was visualised by the iBright imaging system (ThermoFisher). All other genes were visualised with standard UV imaging. Bands were quantified by Fiji (85).

**Table 1:**
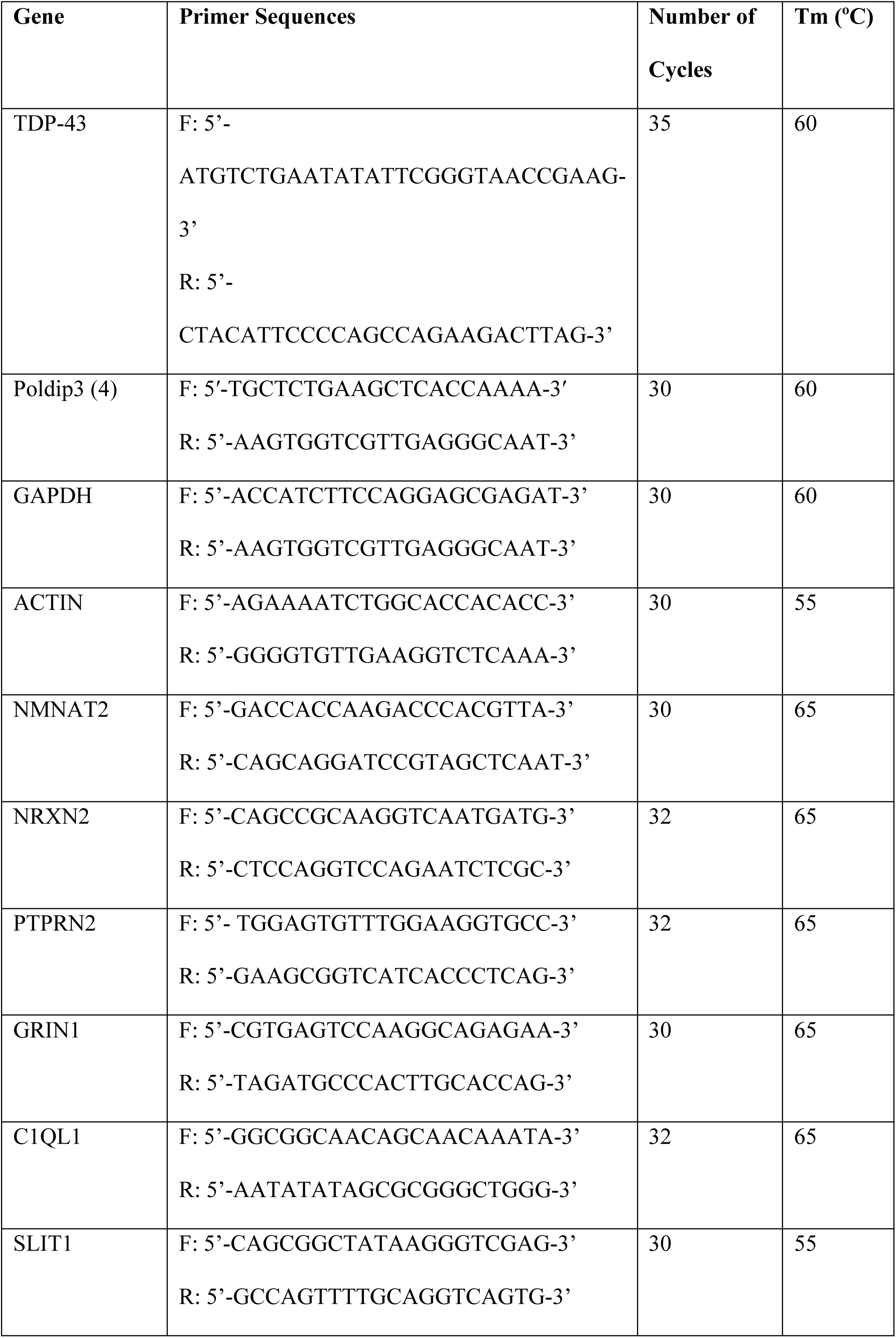
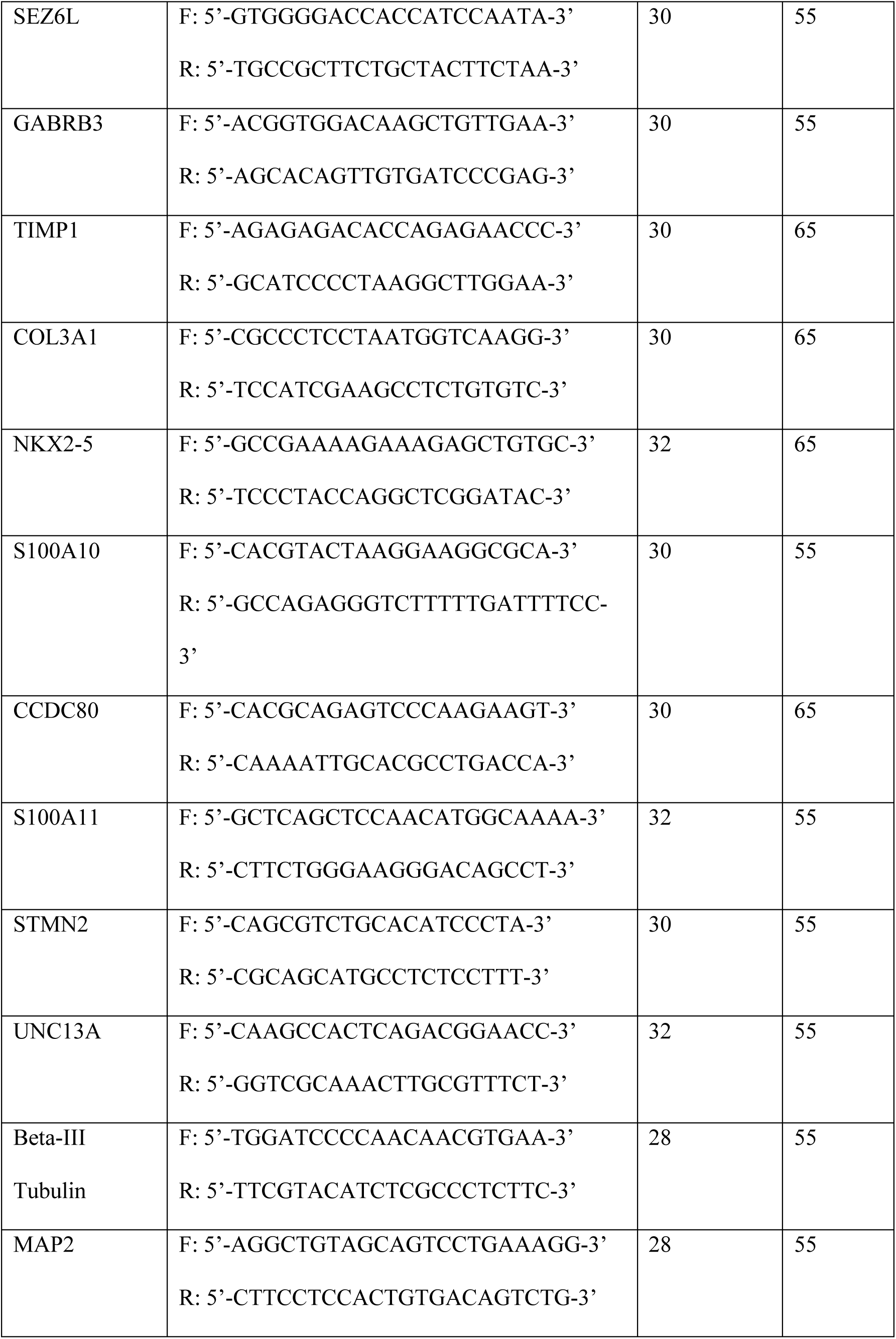
Primers used for RT-PCR experiments. F = Forward primer, R = Reverse primer.

### RNA-Sequencing

Next-generation RNA sequencing (RNA-Seq) was carried out on three independent RNA extractions per CRISPR clones and wild type (WT) SH-SY5Y cells as control. After RNA extraction, samples with A260/280 reads between 1.8 and 2.1 and RNA integrity numbers (RIN) of 10, determined by nanodrop 2000 (Thermo Fisher) and Agilent 2100 BioAnalyser (Agilent), respectively, were used for library preparation. Libraries were generated from >2µg of total RNA following poly(A) selection by Genewiz (Azenta Life Sciences). Sequencing using NovaSeq 6000 (Illumina) generated 2×150-bp reads at a minimum of 30 million read pairs per sample.

### Gene expression profiling

All sequencing reads passed all standard quality metrics as summarised by FastQC v0.12.1 (86). Reads were pseudo-aligned to Gencode v44 transcripts using kallisto v0.48 (87). Gene-level and transcript-level transcript per million (TPMs) were generated from the kallisto abundance files using the R package tximport v1.24.0 (88). Reads were projected to the Gencode v44 transcripts fasta file using kallisto pseudobam and consensus sequences derived using SAMtools v1.20 (89). Potential off-target guide locations were identified using BLAST+ v2.14.1 “blastn-short” (90), allowing up to 4 mismatches, and manually inspected for relevant mutations across all sequenced samples, identifying no off-target exonic mutations. The potential for intronic or intergenic off-target mutations was determined by comparing genomic locations with up to 4 mismatches from the CRISPR guides to the TPM and differential expression analysis data for the containing (intronic) or nearest downstream (intergenic) gene. Univariate differential expression analysis was performed on the gene-level and transcript-level using sleuth v0.30.0 (91) in R v4.2.1 (92). The thresholds for significance were q < 0.05 or 0.1 and absolute log2(fold change) ≥1. Volcano plots were generated on GraphPad Prism version 9.4.1 for Windows. Gene Set Enrichment Analysis (GSEA) was run using the R package fgsea (93) using the prerank setting based on πvalues (-log10(q- value) x log2(fold change)) (94). Enriched sets were identified from the v2023.2 MSigDB gene ontology biological processes and cellular components gene sets (95). Gene ontology information was presented using SRPlot (96). Isoform switching analysis was performed based on the kallisto abundance files using the IsoformSwitchAnalyzeR v1.16.0 (97). The thresholds for significance were q < 0.05 and absolute difference in isoform fraction (dIF) ≥0.1.

### Antibodies

The antibodies used in this study include mouse beta-III tubulin (1:200, Antibodies.com, A86691), chicken MAP2 (1:5000, Antibodies.com, A85363), rabbit recombinant anti-TDP-43 (1:1000; Proteintech, 80002-1-RR), rabbit cleaved caspase-3 (1:1000; Cell Signalling, 9664S), rabbit anti-TDP-43 antibody (1:2000, Proteintech, 10782-2-AP) and a rabbit anti-GAPDH antibody (1:1000, Cell Signalling, #2118S), goat anti-rabbit IgG (H+L) DyLight 488 secondary (1:500, Invitrogen), goat anti-mouse IgG (H+L) DyLight 550 secondary (1:500, Invitrogen), goat anti-rabbit IgG (H+L) DyLight 550 secondary (1:500; Invitrogen), goat anti-chicken IgY (H+L) Alexa Fluor 650 secondary (1:500, Invitrogen) and goat anti-rabbit IgG (H+L), DyLight 680 (1:10,000; Invitrogen).

### SH-SY5Y differentiation

SH-SY5Y cells were plated on collagen-coated (0.05mg/ml, Sigma) plates in standard media (DMEM/F12 +10% FBS) for 24 hours before media was replaced with DMEM + 15% FBS containing 10µM retinoic acid (Sigma, R2625). This media was refreshed every 2 days. At day 5, cells were washed three times with serum free DMEM, before continued culturing in serum free DMEM containing 50ng/ml BDNF (Qkine, Qk050). Medium was refreshed every 2 days and cells harvested or fixed after 4 days in BDNF for further analysis.

### Solubility fractionation and western blot analysis

The fractionation for protein solubility was conducted following a protocol described in Chen et al. (98). Cells were lysed in RIPA buffer (150 mM NaCl, 1% NP-40, 0.5% sodium deoxycholate, 0.1% SDS, 50 mM Tris pH 8.0 and protease and phosphatase inhibitors), sonicated for 10 seconds and centrifuged at 12,000rpm for 20 min at 4°C. Following centrifugation, the supernatant was collected as the soluble RIPA fraction. The pellet was washed once with RIPA buffer and then suspended in 10uL Urea buffer (7 M Urea, 2 M thiourea, 4% CHAPS and 30 mM Tris pH 8.5) as the insoluble, detergent-resistant fraction. Protein concentration was determined by DC Protein Assay (BioRad). 10ug of lysate of the RIPA fraction and the equivalent volume of the urea fraction were loaded for SDS-PAGE.

Protein quantification and western blotting were done as previously described (98). Western blot quantification were performed using Fiji. Integrated band intensities were normalised to that of the loading control.

### Immunofluorescence

Cells were fixed in 4% paraformaldehyde for 30 min and washed with PBS three times for 5 min. Cells were permeabilised by incubation in 0.5% Triton X-100 (Sigma) in PBS for 15 min at room temperature, followed by blocking in 1% goat serum(diluted in PBS) for 1 hour at room temperature. Cells were incubated with primary antibodies diluted in blocking solution overnight at 4°C. After washing in PBS three times for 5 min, cells were subsequently incubated with fluorescent secondary antibodies diluted in the blocking solution for 1 hour at room temperature. DAPI (Sigma) was then used to stain for nuclei before being mounted on coverslips using FluorSave Reagent (Merck).

### Neurite growth analysis

Cells stained with MAP2 were imaged on Axio Observer 7 (Zeiss). Neurites from individual cells were traced with NeuronJ plugin on Fiji (99) and quantified for sum neurite length (µm). A minimum of thirty cells were traced across three individual repeats.

### Data Availability

RNA-Sequencing data for all genotypes is now available following the SRA accession number: PRJNA1191337.

## Acknowledgements

We thank Dr Graham Cocks, head of Genome Editing and Embryology Core, King’s College London, for professional advice and support for establishing CRISPR-knock-in cells for the project.

This work was supported by the Academy of Medical Science (AMS, with Wellcome Trust, BEIS, the British Heart Foundation and Diabetes UK) [SBF006\1088] to HJC; and the Biology departmental PhD studentship to MM. RTG and ASM were supported by ASM’s York Against Cancer (York, UK) 30th Anniversary Lectureship in Cancer Informatics.

## Conflict of Interest Statement

We have no conflicts of interest.

## Abbreviations

ALS: Amyotrophic Lateral Sclerosis
FTD: Frontotemporal Dementia
AD: Alzheimer’s Disease
TDP-43: TAR DNA/RNA-Binding Protein 43

## Notes

### Competing Interest Statement

The authors have declared no competing interest.

### Summary of Updates

We noticed an effor of misplacing a K263E RNA-sequence sample, corrected it and re-do all K263E analysis (Supplementary data 1, supplementary figure 2, 4 and 6). Further analysis and new figures were included in this revision to rule out CRISPR off-target editing impacting on our results (Supplementary figure 3). Finally, a new correlation plot is now included to compare our homozygous TDP-43 mutant cells with the published si-TDP-43 cells (Supplementary figure 4B).

