## Supplementary figure for "Familial ALS/FTD-associated RNA-Binding deficient TDP-43 mutants cause neuronal and synaptic transcript dysregulation *in vitro*"

### Supplementary Figure 1

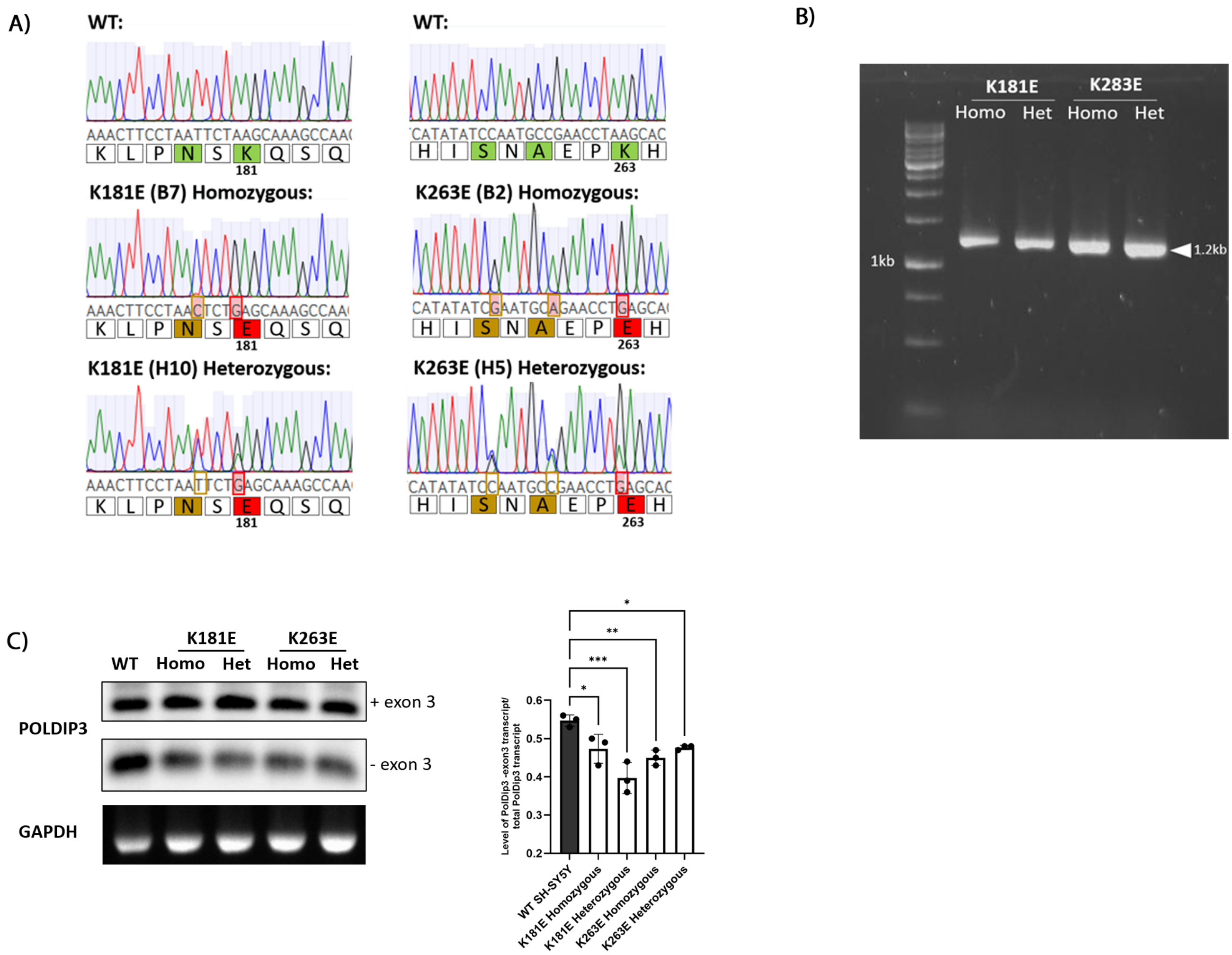

### Supplementary Figure 2

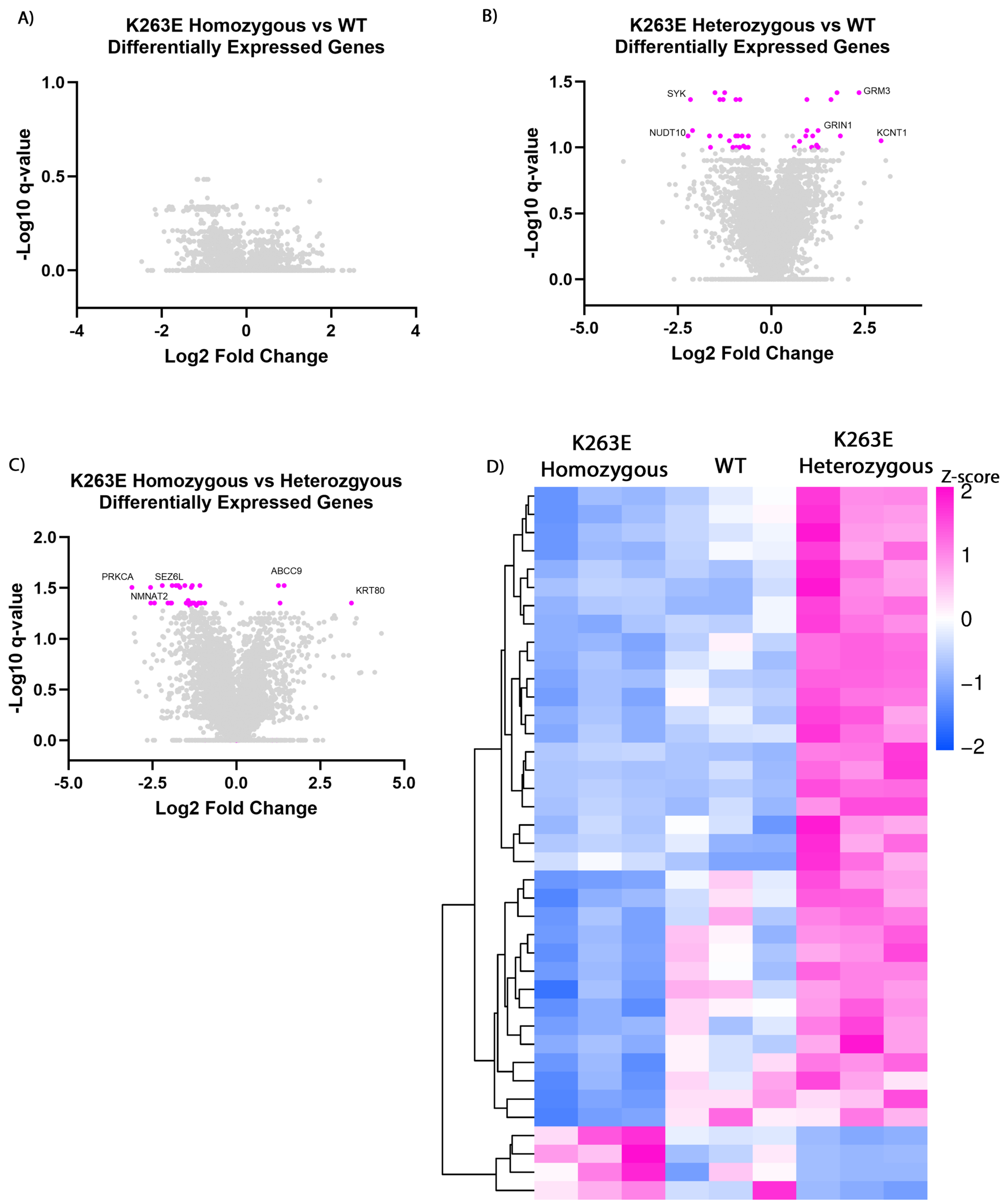

### Supplementary Figure 3

A) WGE ID: #1188620536

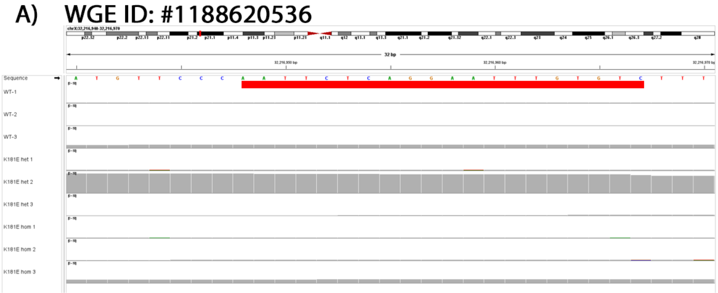

B) WGE ID: #1117723179

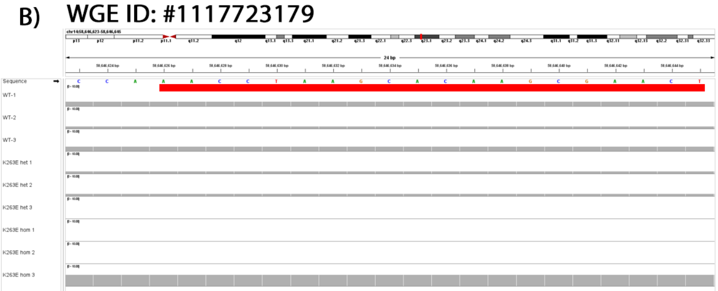

C) WGE ID: #1150237452

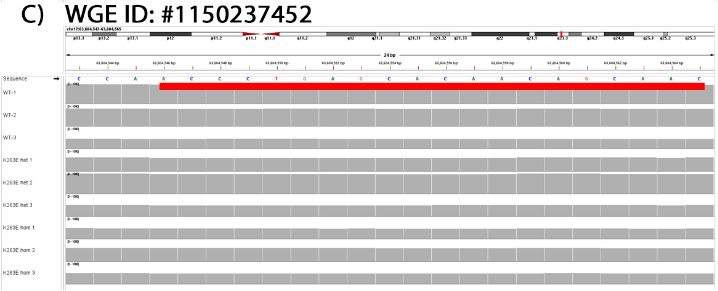

D) WGE ID: #1100984314 #1129748034 #1056910716

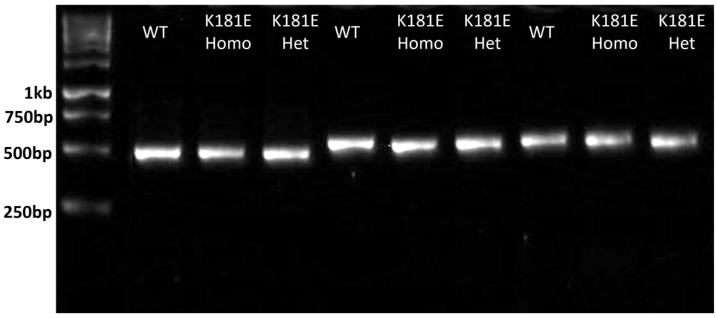


E) WGE ID: #1109273067 #948477925 #1024616983

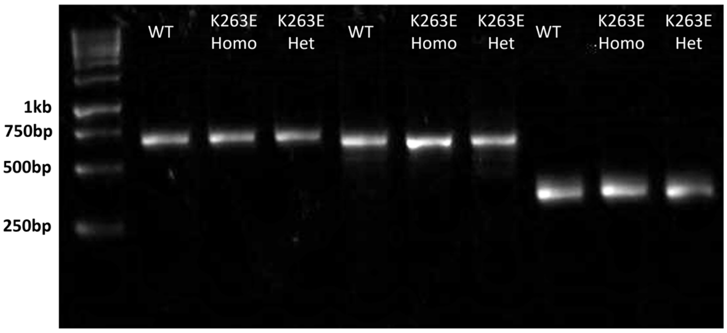


F)

**#1100984314**

| | KO | $R^2$ | INDEL | % |
|---|---|---|---|---|
| WT | 0.0 | 0.99 | 0 | 100.0 |
| K181E Homo | 0.0 | 0.99 | 0 | 100.0 |
| K181E Het | 0.0 | 0.98 | 0 | 100.0 |

**#1129748034**

| | KO | $R^2$ | INDEL | % |
|---|---|---|---|---|
| WT | 0.0 | 1.00 | 0 | 100.0 |
| K181E Homo | 0.0 | 1.00 | 0 | 100.0 |
| K181E Het | 0.0 | 1.00 | 0 | 100.0 |

**#1056910716**

| | KO | $R^2$ | INDEL | % |
|---|---|---|---|---|
| WT | 0.0 | 1.00 | 0 | 100.0 |
| K181E Homo | 0.0 | 1.00 | 0 | 100.0 |
| K181E Het | 0.0 | 1.00 | 0 | 100.0 |

G)

**#1109273067**

| | KO | $R^2$ | INDEL | % |
|---|---|---|---|---|
| WT | 0.0 | 0.98 | 0 | 100.0 |
| K263E Homo | 0.0 | 0.98 | 0 | 100.0 |
| K263E Het | 0.0 | 0.98 | 0 | 100.0 |

**#948477925**

| | KO | $R^2$ | INDEL | % |
|---|---|---|---|---|
| WT | 0.0 | 0.98 | 0 | 100.0 |
| K263E Homo | 0.0 | 0.98 | 0 | 100.0 |
| K263E Het | 0.0 | 0.98 | 0 | 100.0 |

**#1024616983**

| | KO | $R^2$ | INDEL | % |
|---|---|---|---|---|
| WT | 0.0 | 1.00 | 0 | 100.0 |
| K263E Homo | 0.0 | 1.00 | 0 | 100.0 |
| K263E Het | 0.0 | 1.00 | 0 | 100.0 |

#### Supplementary Figure 4

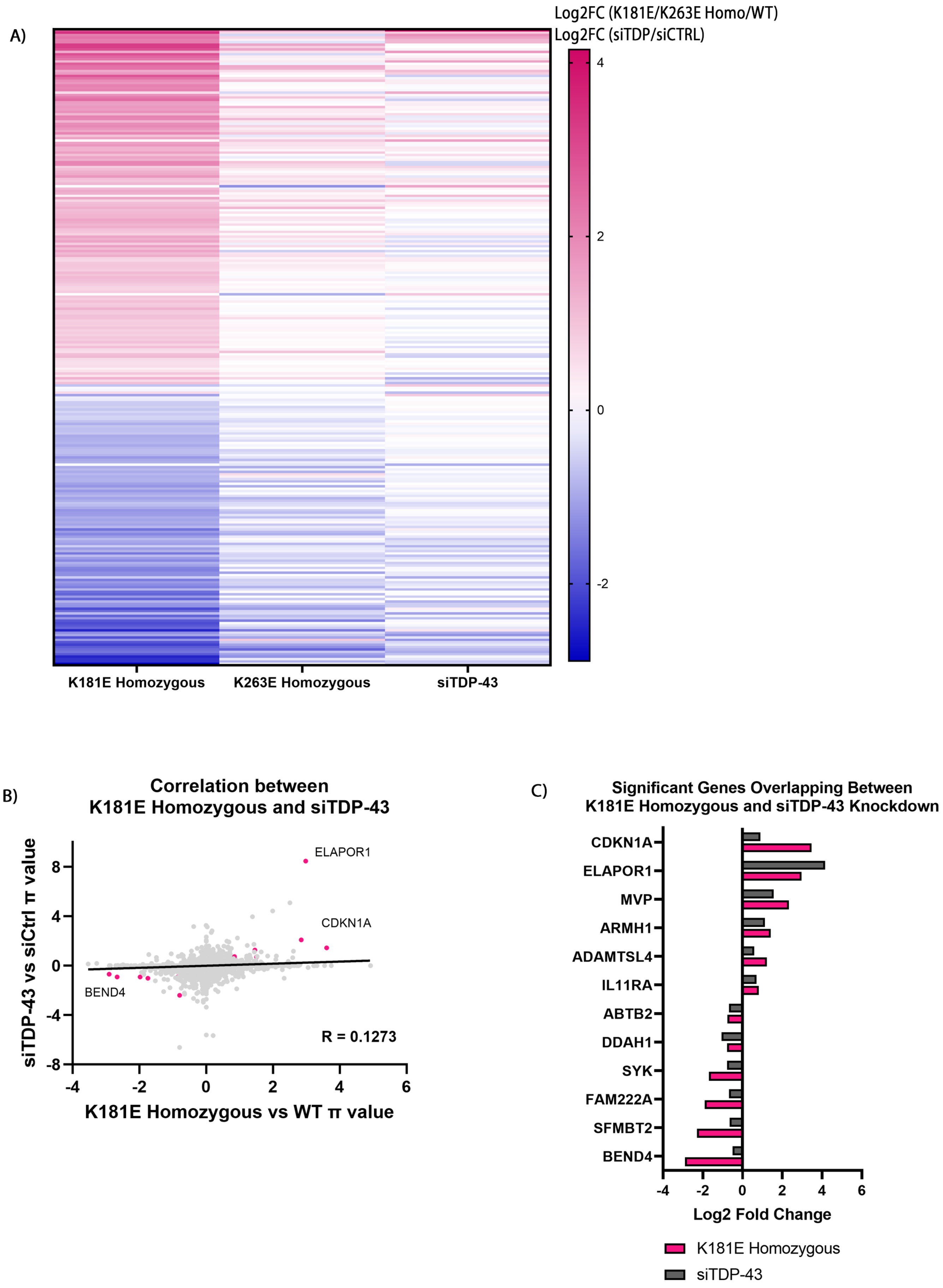

### Supplementary Figure 5

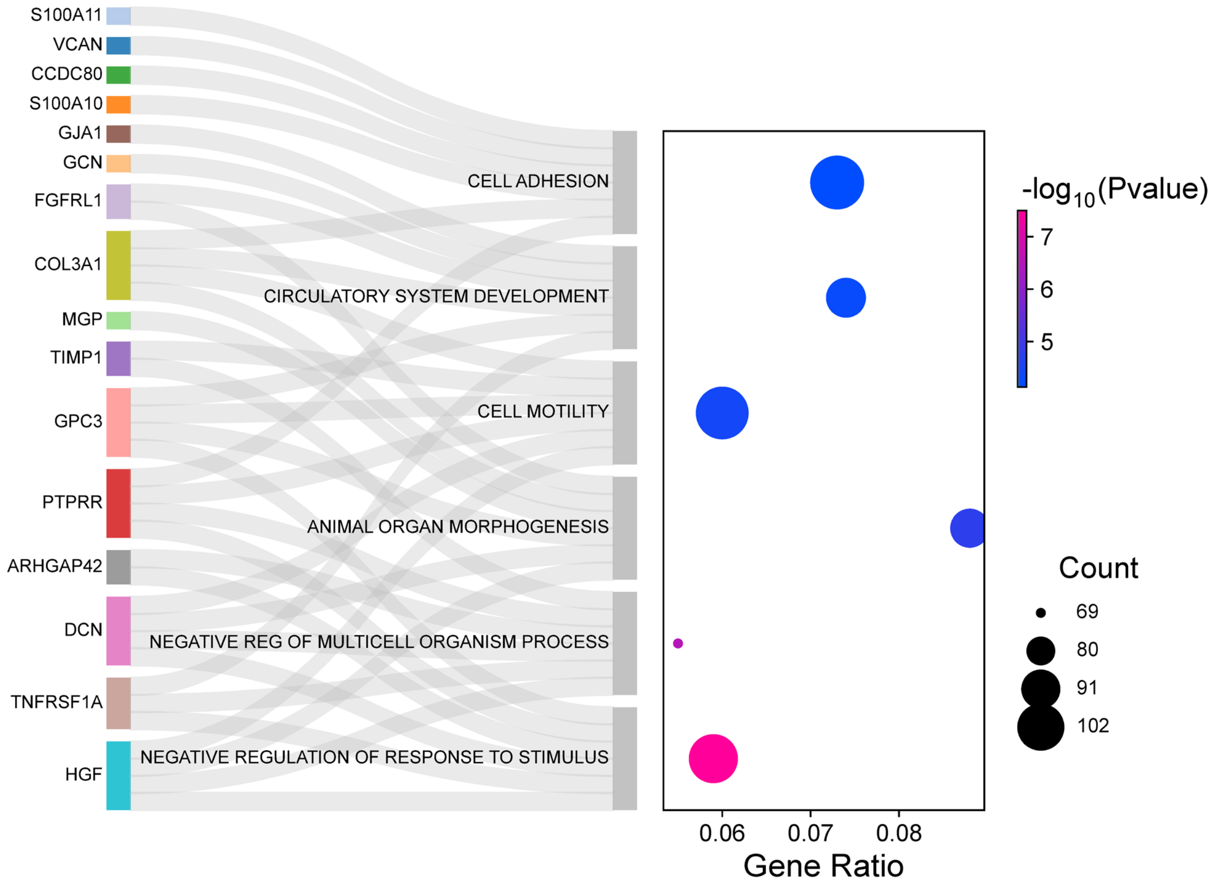


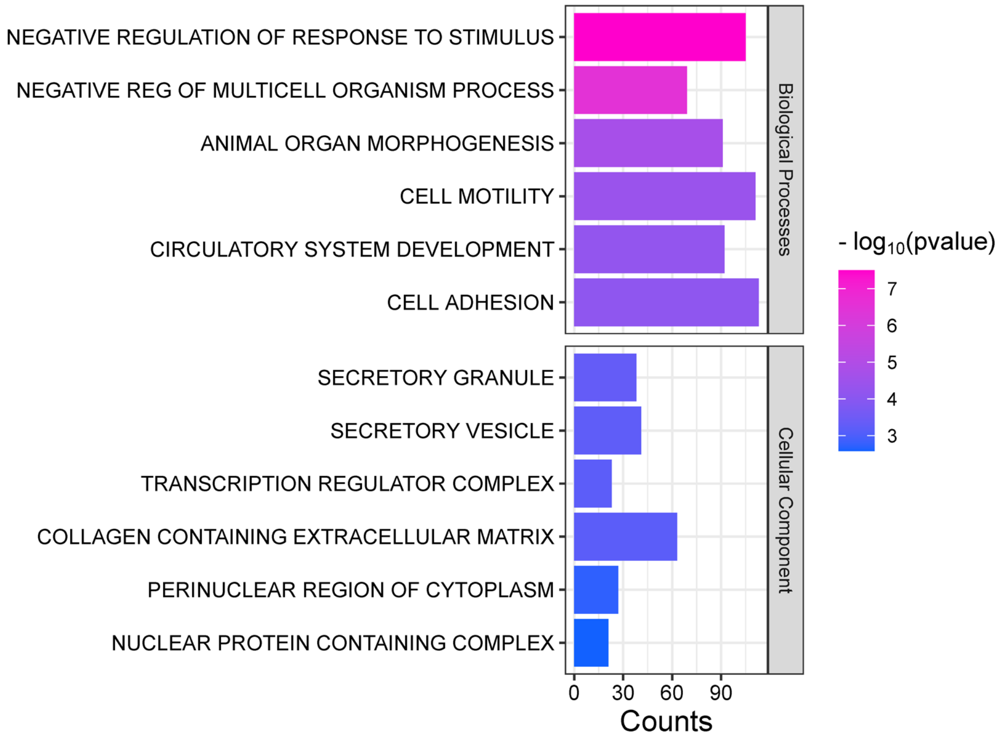

### Supplementary Figure 6

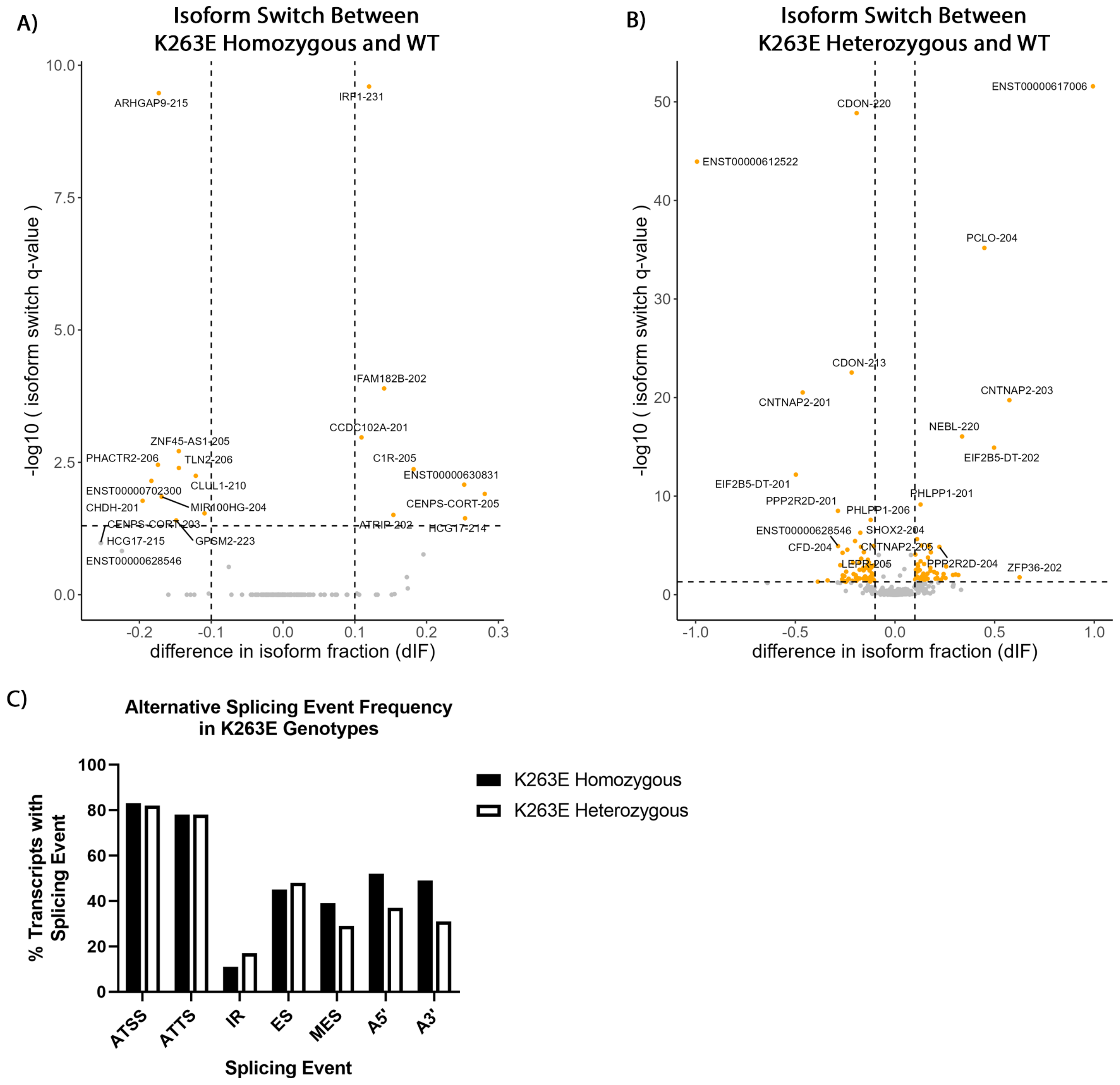

### Supplementary Figure 7

A)

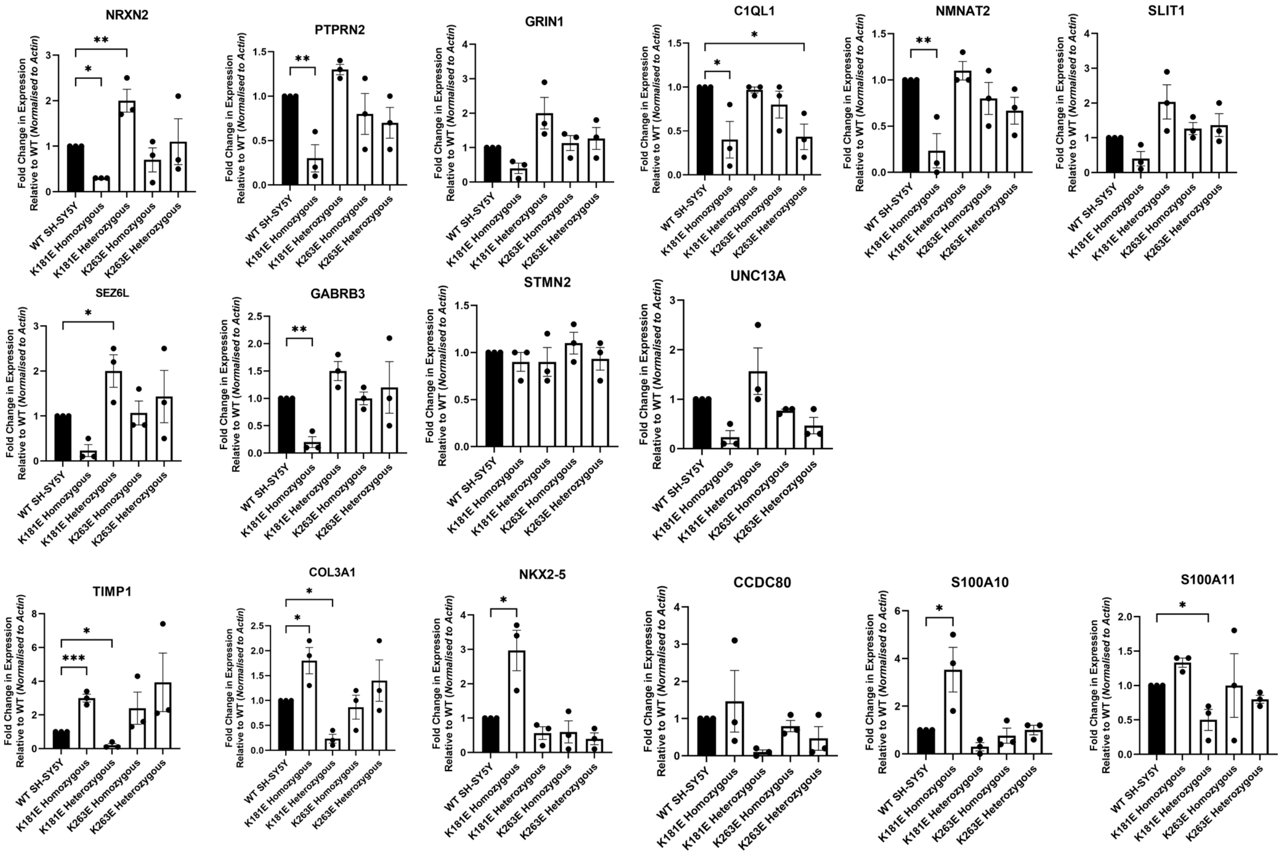


B)

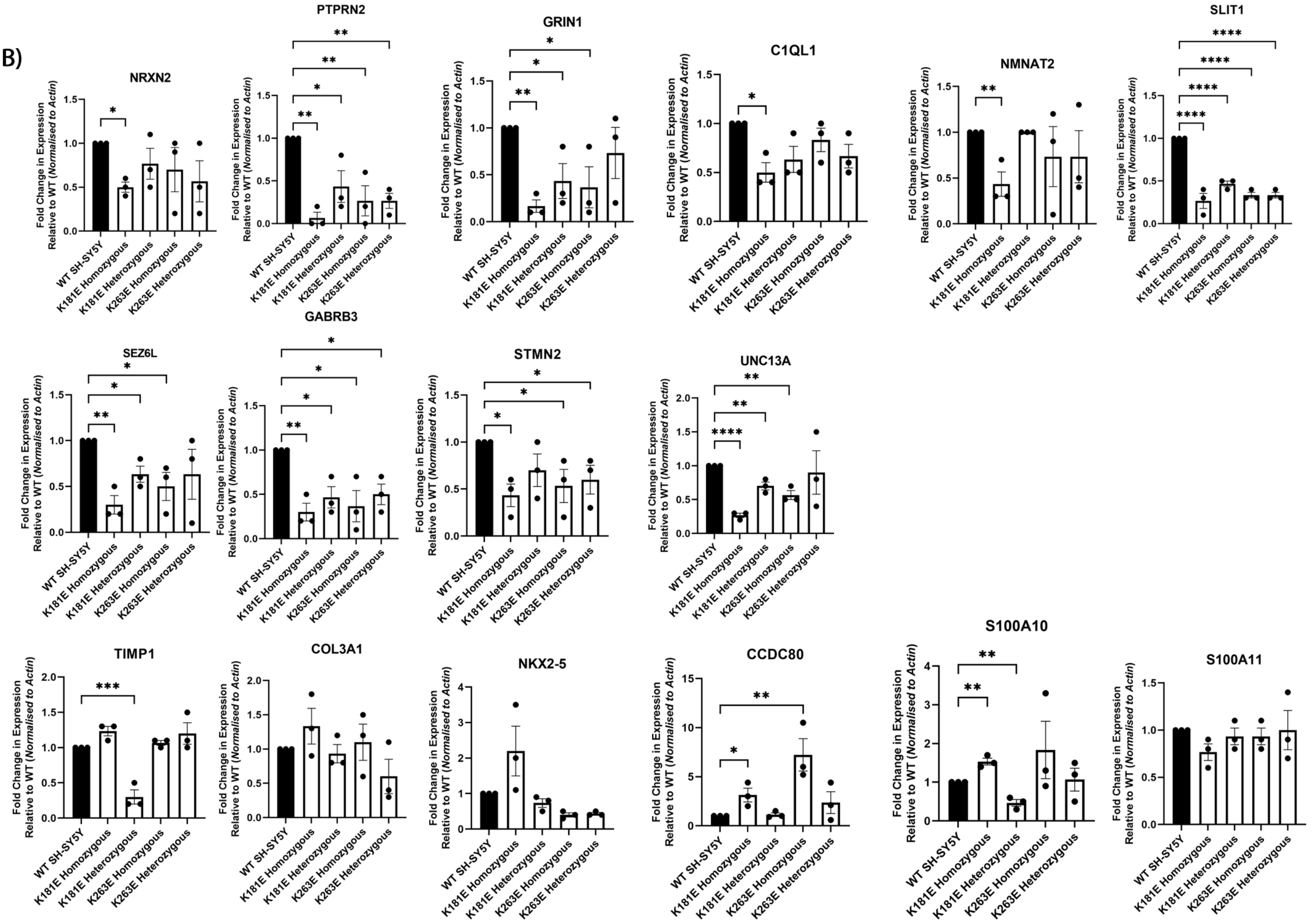

### Supplementary Figure 8

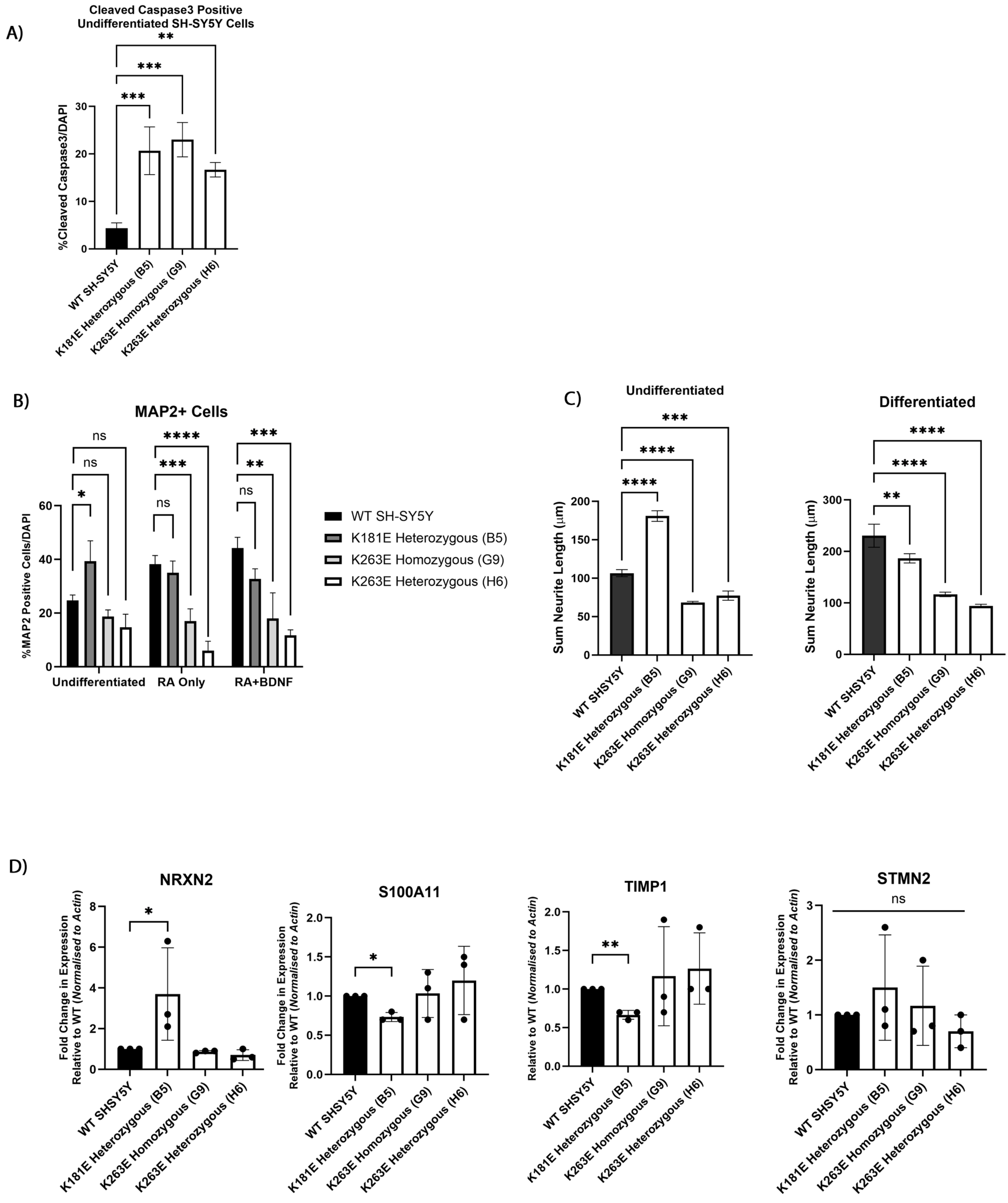
