## Supplementary figure legend for "Familial ALS/FTD-associated RNA-Binding deficient TDP-43 mutants cause neuronal and synaptic transcript dysregulation *in vitro*"

**Supplementary Figure 1: Validation of CRISPR/Cas9-edited K181E and K263E-TDP-43 cell lines.** **A.** Sanger sequencing chromatogram showing CRISPR/Cas9-introduced nucleotide substitutions in the targeted genomic area. Gold box indicates silent mutations that were implemented in the HDR template to prevent repetitive Cas9 cutting, the red box indicates Lysine to Glutamic acid substitutions (K181E or K263E). **B.** Examination of the intactness of TDP-43 cDNA using RT-PCR. The single clean band at the expected size of 1.2kb suggests that there is no CRISPR/Cas9-induced prominent insertion or deletion in TDP-43 transcript. **C.** *PolDip3* mRNA splicing assay of unmodified WT and edited TDP-43 cells. *GAPDH* was used as the loading control. The levels of PolDip3 long and short transcripts from three independent experiments were quantified and normalised to GAPDH. Means and SDs are shown**.** The data was analysed by One-way ANOVA followed by Dunnet’s post-test. * p<0.05, ** p<0.01, *** p<0.001.

**Supplementary Figure 2: No significant transcriptomic profile change in K263E-TDP-43 cells. A-B.** Volcano plots of transcriptomic profile of K263E-TDP-43 homozygous (A) or heterozygous cells (B) relative to un-edited control SH-SY5Y cells. No gene reaches a significant adjusted *p* value (-Log10(q<0.1)), indicating no significant DEGs in either K263E-TDP-43 cell lines compared to WT.

**Supplementary Figure 3: No evidence for off-target CRISPR modifications. A-C.** Confirming the absence of exonic off-target modification in the RNA-seq sample results. The sequencing of all conditions were investigated around the putative off-target sites. No evidence for *DMD* (WGE ID#1188620536) (**A**), *DACT1* (#1117723179) (**B**) or *TACO1* (#1150237452) (**C**) off-target modification. guideRNA similarity site highlighted with red box. **D-E.** PCR of top 3 K181E (**D**) and K263E (**E**) intronic/intergenic off-targets (WGE IDs) for unedited WT cells, homozygous and heterozygous cells. **F-G.** Presence of in/dels detected by DECODR (99) for K181E (**F**) and K263E (**G**). KO = Knock-out, R^2^ = how well the sequence fits the template using Pearson’s coefficient correlation, INDEL = insertion/deletion number.

**Supplementary Figure 4: Comparison of gene expression changes in K181E-TDP-43 homozygous, K263E-TDP-43 homozygous and published TDP-43 knockdown cells. A.** Heat map comparing the fold changes in gene expression in homozygous K181E or K263-TDP-43 CRISPR cells or in TDP-43 knockdown cells. Analysed genes were those with an adjusted *p* value (q) < 0.1 identified in K181E-TDP-43 homozygous cells. B. Scatter plot and linear model relating π-values (-log10(q-value) x Log2FC) of K181E-TDP-43 homozygous vs WT, and siTDP-43 vs siCtrl. Significant DEGs shared between both datasets are highlighted in pink. π-values were weakly correlated (Pearson correlation coefficient R = 0.1273). **C.** The fold changes of the 12 DEGs shared in our homozygous K181E-TDP-43 dataset and Melamed’s siTDP-43 dataset.

**Supplementary Figure 5: Gene set enrichment analysis highlighted a diverse set of pathways from the up-regulated genes in K181E homozygous cells.** Gene set enrichment analysis for DEGs up-regulated in K181E-TDP-43 homozygous cells (compared to K181E-TDP-43 heterozygous), using gene ontology biological processes and cellular components gene sets from MSigDB. πvalues (-log10(p-value) x log2(fold change)) were used as the rank of significance. The most significant gene ontology terms for biological processes and cellular components and associated gene are shown with associated significance (-log10(Pvalue), shown in colour scale). Gene ratio= (Gene count from dataset associated with pathway/ Total gene count within pathway dataset).

**Supplementary Figure 6: Isoform switch analysis of K263E-TDP-43 cells. A-B.** Volcano plots generated from IsoformSwitchAnalyzeR showing significant isoform switches in K263E-TDP-43 homozygous cells (A) or heterozygous cells (B) relative to un-edited control SH-SY5Y cells. -Log10(q<0.05) plotted against difference in isoform fraction**.** Significant changes are recognised by an absolute difference in isoform fraction of 0.1, highlighted in yellow. **C.** Alternative splicing events caused by K263E-TDP-43 expression. The alternative splicing events are identified by IsoformSwitchAnalyzeR, analysing all transcript isoforms identified as significantly changed in (A) and (B). ATSS, Alternative Transcript Start Site; ATTS, Alternative Transcript Termination Site; IR, Intron Retention; ES, Exon Skipping; MES, Mutually Exclusive Exons; A5’, Alternative 5’ Splicing; A3’, Alternative 3’ Splicing.

**Supplementary Figure 7: RT-PCR validation of selected DEGs in K181E-TDP-43 cells in undifferentiated and differentiated stages.** Quantification of RT-PCR of selected DEGs and known TDP-43 targets (STMN2 and UNC13A) in undifferentiated (A) and differentiated cells (B). The data were obtained from three independent extractions. Means and SDs are shown**.** The data was analysed by One-way ANOVA followed by Dunnet’s post-test. Only significant changes are highlighted. * p<0.05, ** p<0.01, *** p<0.001, **** p<0.0001.

**Supplementary Figure 8: The transcriptomic and cellular disturbances are validated in independent CRISPR lines. A.** Reduced cell viability in TDP-43 cells. Cells were stained for cleaved caspase-3, the percentages of which are quantified from three independent experiments. Means and SDs are shown**.** The data was analysed by One-way ANOVA followed by Dunnet’s post-test, showing K181E or K263E-TDP-43 cells are less viable than unmodified control. **B.** Impaired differentiation efficiency in TDP-43 cells. Cells were stained for MAP2, marker of neuronal differentiation and the percentages of MAP2-positive cells at different stages were quantified from three independent studies. Means and SDs are shown. The data were analysed by One-way ANOVA followed by Dunnet’s post-test. **C.** Sum neurite length from MAP2 staining was quantified for individual cells across three independent experiments for the individual clones. Means and SDs are shown. The data were analysed by One-way ANOVA followed by Dunnet’s post-test. **D.** Four genes were selected for RT-PCR analysis in undifferentiated cells. The levels of selected genes from three independent studies are quantified. Means and SDs are shown**.** The data was analysed by One-way ANOVA followed by Dunnet’s post-test. * p<0.05, ** p<0.01, *** p<0.001, **** p<0.0001.
