## Supplementary table 1 for "Familial ALS/FTD-associated RNA-Binding deficient TDP-43 mutants cause neuronal and synaptic transcript dysregulation *in vitro*"

**Supplementary Table 1: Primers used for off-target PCR and sequencing.** F = forward primer. R = reverse primer.

| **WGE ID** | **Primer Sequences** | **Number of Cycles** | **Tm (^o^C)** |
| --- | --- | --- | --- |
| #1100984314 PCR | F: 5’- AAGGGCTGTGAAGGAGAAGC -3’  R: 5’- CAGTGGCTTTGGCCTGTAGA -3’ | 35 | 60 |
| #1100984314  Sequencing | F: 5’- GGCCTGTCGGGAAGACAATACATC-3’  R: 5’- GCGCCTATCTGCATTCTCTG-3’ | N/A | N/A |
| #1129748034  PCR | F: 5′-AGAAATACAGAAACTACAGGGCTAT -3′  R: 5’- AAGGGCCATCGGTGTATGAC-3’ | 35 | 60 |
| #1129748034  Sequencing | F: 5’- AGGGCCTGTTGGGAAGACATAC-3’  R: 5’- GGCCATGTCTCCGGTTGTATTTAG-3’ | N/A | N/A |
| #1056910716  PCR | F: 5’ CACCTCCATCATAGGACGCC -3’  R: 5’- TGTGCTGATGATGCTCAGGG -3’ | 35 | 60 |
| #1056910716  Sequencing | F: 5’- CAGCAATGCAGTGTTAATTGCTAC-3’  R: 5’- CCCATCCAGTAGCAGAAAGCTC-3’ | N/A | N/A |
| #1109273067  PCR | F: 5’- TTTAGTAGTGTTGGGTCGCACA -3’  R: 5’- CCCCAACTGCTTTGTAGCAC -3’ | 35 | 60 |
| #1109273067  Sequencing | F: 5’- GAAGTGGTGGATGTCGTCATCCTC-3’  R: 5’- CACCTGGATTACCACCAAATCTTC-3’ | N/A | N/A |
| #948477925  PCR | F: 5’- AGACGTTCTTATGGTGCGGC -3’  R: 5’- GGATGCTGATCCCCAACCAA -3’ | 35 | 60 |
| #948477925  Sequencing | F: 5’- GGATGTGATGGATGTCTTCGTCCC-3’  R: 5’- GCTGAACGCACCAAAGTTTATC-3’ | N/A | N/A |
| #1024616983  PCR | F: 5’- CCCCCTCTGCCAAGTGTTAC-3’  R: 5’- AACTGGTCCCATGATGCAGG-3’ | 35 | 60 |
| #1024616983  Sequencing | F: 5’- AATGAAGATAACAGAACCTGCC-3’ | N/A | N/A |
